# Nuclear-enriched protein phosphatase 4 ensures outer kinetochore assembly prior to nuclear dissolution

**DOI:** 10.1101/2022.09.01.505886

**Authors:** Helder Rocha, Patrícia A. Simões, Jacqueline Budrewicz, Pablo Lara-Gonzalez, Ana Xavier Carvalho, Julien Dumont, Arshad Desai, Reto Gassmann

**Affiliations:** Instituto de Investigação e Inovação em Saúde - i3S, Universidade do Porto, Porto, Portugal; Ludwig Institute for Cancer Research, San Diego Branch, La Jolla, USA; Division of Biological Sciences, Department of Cellular and Molecular Medicine, University of California San Diego, La Jolla, USA; Department of Molecular and Medical Genetics, Oregon Health & Science University (OHSU), Portland, Oregon, USA; Division of Reproductive & Developmental Sciences, Oregon National Primate Research Center (ONPRC), Beaverton, Oregon, USA; Department of Developmental and Cell Biology, School of Biological Sciences, University of California, Irvine, USA; Institut Jacques Monod, CNRS, UMR 7592, University Paris Diderot, Sorbonne Paris Cité, Paris, France

## Abstract

A landmark event in the transition from interphase to mitosis in metazoans is nuclear envelope breakdown (NEBD). Many events important for mitosis occur prior to NEBD, including condensation of replicated chromosomes and assembly of kinetochores to rapidly engage spindle microtubules. Here we show that nuclear-enriched protein phosphatase 4 (PP4) ensures robust assembly of the microtubule-coupling outer kinetochore prior to NEBD. In the absence of PP4, chromosomes exhibit extended monopolar orientation after NEBD and subsequently mis-segregate. A secondary consequence of diminished outer kinetochore assembly is defective sister chromatid resolution. After NEBD, a cytoplasmic activity compensates for PP4 loss, leading to outer kinetochore assembly and recovery of chromosomes from monopolar orientation to significant biorientation. The Ndc80-Ska microtubule-binding module of the outer kinetochore is required for this recovery. PP4 associates with the inner kinetochore protein CENP-C; however, disrupting the PP4–CENP-C interaction does not perturb chromosome segregation. These results establish that PP4-dependent outer kinetochore assembly prior to NEBD is critical for timely and proper engagement of chromosomes with spindle microtubules.

## INTRODUCTION

Successful metazoan mitosis requires that replicated chromosomes are extensively re-organized prior to NEBD so that they can be segregated to daughter cells by the microtubule-based spindle. This entails physical compaction and resolution of sister chromatids into discrete rod-shaped structures (chromosome condensation) and assembly of a microtubule attachment site at the centromeric locus on each sister chromatid (the kinetochore) (Batty and Gerlich, 2019; Musacchio and Desai, 2017; Navarro and Cheeseman, 2021; Paulson *et al*., 2021). In late prophase the chromatin-proximal inner kinetochore recruits the microtubule-coupling outer kinetochore, allowing rapid engagement of spindle microtubules after NEBD. Prometaphase chromosomes typically establish initial attachments to one spindle pole (mono-orientation) before forming stable load-bearing attachments that correctly connect sister kinetochores to opposite poles (bi-orientation). Bi-orientation is accompanied by chromosome alignment at the spindle equator, and a mitotic checkpoint ensures all chromosomes have time to bi-orient before sister chromatids separate in anaphase (Lara-Gonzalez *et al*., 2021). Efficient bi-orientation and alignment of chromosomes is crucial in the rapidly dividing cells of the early embryo, which lack a robust checkpoint response (Duro and Nilsson, 2020).

Chromosome condensation, kinetochore assembly, and kinetochore-microtubule interactions are extensively regulated by kinase and phosphatase activities. Regulation by Ser/Thr protein phosphatases (PP) is best understood for PP1 and PP2A (Moura and Conde, 2019; Nilsson, 2018), whereas the PP2A-like PP4 has received comparatively little attention (Park and Lee, 2020). In its most common form, PP4 is a heterotrimer consisting of the catalytic subunit PP4c and two regulatory subunits, PP4R2 and PP4R3, the latter targeting PP4 to substrates in the nucleus and cytoplasm through its N-terminal EVH1 domain, which binds to FxxP or MxPP motifs (Ueki *et al*., 2019; Karman *et al*., 2020).

In dividing cells PP4 acts on both chromosomes and the microtubule cytoskeleton. Binding of human PP4 to the *wings apart-like* (Wapl) protein promotes cohesin release during prophase (Ueki *et al*., 2019), and targeting of *D. melanogaster* PP4 to the inner kinetochore protein CENP-C is required for mitotic centromere integrity (Lipinszki *et al*., 2015; Torras-Llort *et al*., 2020). The *C. elegans* PP4c homolog PPH-4.1 promotes chromosome pairing and synapsis during female meiotic prophase and microtubule severing activity in mature oocytes (Gomes *et al*., 2013; Guo *et al*., 2022; Han *et al*., 2009; Sato-Carlton *et al*., 2014). Human and *D. melanogaster* PP4 localizes to mitotic spindle poles, and PP4c inhibition produces aberrant spindles with decreased centrosomal γ-tubulin levels (Brewis *et al*., 1993; Hastie *et al*., 2000; Helps *et al*., 1998; Martin-Granados *et al*., 2008). Similarly, *C. elegans* PPH-4.1 was reported to be required for mitotic centrosome maturation in the early embryo (Sumiyoshi *et al*., 2002), while no function has been described for its paralog PPH-4.2.

Here we report that inhibition of the *C. elegans* PP4R3 homolog SMK-1, known so far for post-mitotic functions in longevity and stress resistance (Sen *et al*., 2020; Wolff *et al*., 2006), results in chromosome alignment and segregation defects in the early embryo, which are not attributable to mis-regulation of the microtubule cytoskeleton. Our results establish that the PP4c paralogs PPH-4.1 and PPH-4.2 function redundantly and together with SMK-1 as part of a nuclear-enriched PP4 complex that promotes outer kinetochore assembly prior to NEBD for rapid engagement of spindle microtubules. We show that this novel function of PP4 is crucial for timely and error-free chromosome bi-orientation.

## RESULTS AND DISCUSSION

### PP4 inhibition in the early *C. elegans* embryo results in chromosome congression and segregation defects

In a genome-wide RNAi screen whose hits were enriched for proteins with roles in embryonic mitosis we had identified SMK-1, the sole *C. elegans* PP4 targeting subunit (PP4R3) homolog (Fig. 1A; Fig. S1A) (Rocha *et al*., 2018). To assess whether SMK-1 plays a role in mitosis, we analyzed the effect of its depletion in one-cell embryos co-expressing fluorescent markers for chromosomes and spindle poles/microtubules. One-cell embryos depleted of SMK-1 exhibited a striking mitotic phenotype, with chromosomes scattering across the spindle after NEBD and then traversing back towards the spindle equator prior to anaphase onset (Fig. 1B - F; Movie S1, S2). This phenotype is particularly evident in time-aligned kymographs (Fig. 1C; Fig. S1B) and has not been observed in prior studies focused on the chromosome segregation machinery. Following anaphase onset, a high incidence of chromatin bridges was observed, indicative of persistent chromosome-spindle attachment errors (Fig. 1B, E).

**Figure 1.**
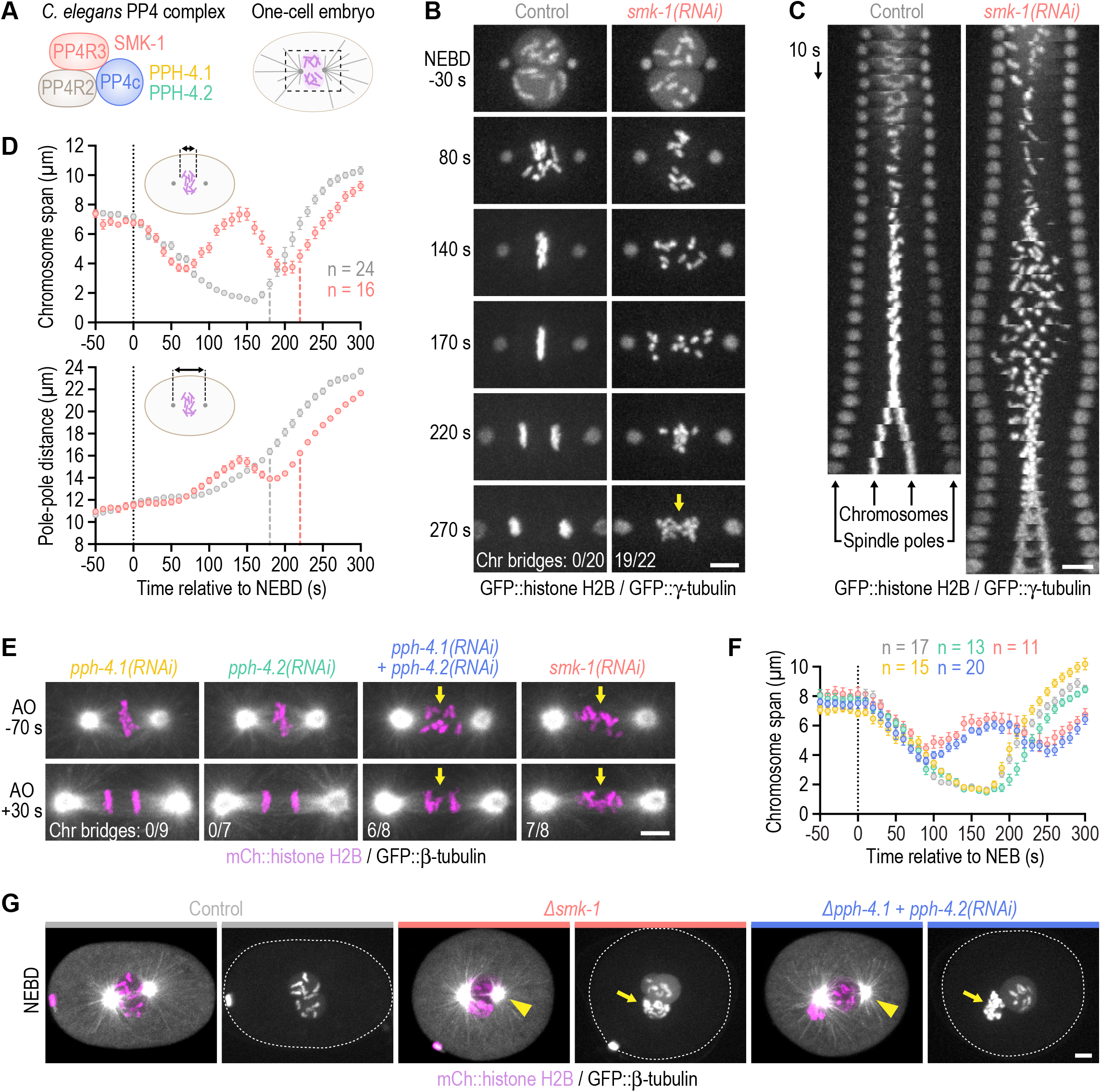
PP4 inhibition in the early *C. elegans* embryo impairs chromosome congression and results in chromosome mis-segregation. **(A)** Cartoon of *C. elegans* PP4 (the catalytic subunit is represented by two paralogs) and of the dividing one-cell embryo with the spindle region imaged in this study outlined by dashed lines. **(B)** Selected images from time-lapse movies of one-cell embryos co-expressing GFP::histone H2B (HIS-58) and GFP::γ-tubulin (TBG-1). Time is relative to NEBD. The number of anaphases with chromatin (chr) bridges, highlighted by the arrow, relative to the total number of anaphases examined is indicated in the last frame. Scale bar, 5 µm. **(C)** Time-aligned kymographs generated from time-lapse movies such as those shown in *(B)* and Movie S1. Scale bar, 5 µm. **(D)** Chromosome span *(top)* and pole-pole distance *(bottom)* versus time relative to NEBD, measured in one-cell embryos such as those shown in *(B)*, as indicated in the schematics. Values correspond to the mean of *n* embryos ± SEM, and vertical dashed lines mark the average time of anaphase onset. Conditions are color-coded as in *(B)*. **(E)** Selected images from time-lapse movies of one-cell embryos co-expressing mCherry::histone H2B (HIS-11) and GFP::β-tubulin (TBB-2). Time is relative to anaphase onset (AO). Arrows highlight defective chromosome congression and segregation in PP4-inhibited embryos. The frequency of anaphase chromatin (chr) bridges is indicated as described for *(B)*. Scale bar, 5 µm. **(F)** Chromosome span versus time, measured in one-cell embryos such as those shown in *(E)* and plotted as described for *(D)*. Conditions are color-coded as in *(B)* - *(E)*. **(G)** Selected images from time-lapse movies of one-cell embryos co-expressing mCherry::histone H2B and GFP::β-tubulin, isolated from controls or homozygous mutant mothers before the onset of sterility. Time point corresponds to NEBD. Arrows point to structurally aberrant extra chromosomes in the female pronucleus, which are carried over into the first mitosis from defective meiotic divisions. This meiosis-derived defect is only observed in the two most penetrant PP4 inhibition conditions shown here. Centrosome maturation remains unaffected by penetrant PP4 inhibition (arrowheads). Scale bar, 5 µm.

To determine whether SMK-1’s role in chromosome segregation reflects its molecular function as the PP4 targeting subunit, we inhibited the PP4 catalytic subunit. Individual inhibition of the PP4c paralogs PPH-4.1 and PPH-4.2 by RNAi or genetic knock-out (*∆pph-4*.*1* and *∆pph-4*.*2*, respectively; Fig. S1A, C - E) had no adverse effect on the first embryonic division. By contrast, co-inhibition of PPH-4.1 and PPH-4.2 phenocopied SMK-1 depletion (Fig. 1E, F; Fig. S1F, G; Movie S3). Thus, PPH-4.1 and PPH-4.2 function redundantly in the early embryo and SMK-1 contributes to mitotic chromosome segregation in the context of PP4 complexes.

Genetic knock-out of *smk-1* (*∆smk-1*; Fig. S1A, C - E) and penetrant inhibition of PP4c produced a variable number of structurally aberrant chromosomes in the maternal pronucleus (Fig. 1G; Fig. S1E), suggesting that PP4 is also required for chromosome segregation during oocyte meiosis. Immunoblotting suggested that this phenotype may not be observed in *smk-1(RNAi)* embryos because of residual protein (Fig. S1H). Thus, PP4c acts during oocyte meiosis, in addition to ensuring proper chromosome segregation in the embryo.

### Chromosomes in PP4-inhibited embryos go through an extended period of mono-orientation before recovering to a bi-oriented state

Inspection of the mitotic PP4 inhibition phenotype revealed unusual chromosome behavior during prometaphase: for the first ~70 s following NEBD chromosomes congressed towards the spindle equator before scattering back towards spindle poles between 70 s and 150 s (Fig. 1B, C). From 150 s until anaphase onset at 230 s (a ~40 s delay relative to controls), chromosomes resumed congression, albeit without achieving full alignment at the spindle equator. The transient scattering of chromosomes manifests as a prominent bounce when the distance between outermost chromosomes along the spindle axis is plotted over time (‘chromosome span’; Fig. 1D, F; Fig. S1G). Plotting the distance between spindle poles over time showed that spindles in PP4-inhibited embryos elongated prematurely during chromosome scattering (Fig. 1D). Spindle length subsequently recovered during chromosome re-congression, and by anaphase onset spindles in PP4-inhibited embryos were the same length as control spindles. The prometaphase bounce in spindle length indicates a delay in the formation of bi-oriented attachments capable of sustaining the tension generated by cortex-localized forces that position the spindle by pulling on astral microtubules (Grill *et al*., 2003; Nguyen-Ngoc *et al*., 2007; Oegema *et al*., 2001; Cheeseman *et al*., 2004). Taken together, these observations suggested that PP4 inhibition results in an extended period of mono-orientation before chromosomes recover to a bi-orientated state. This conclusion was reinforced by the striking consequence of co-inhibiting PP4 and polar ejection forces generated by the chromokinesin KLP-19^Kif4^ (Powers *et al*., 2004): in *klp-19*^*Kif4*^*(RNAi);smk-1(RNAi)* embryos, but not in *klp-19*^*Kif4*^*(RNAi)* or *smk-1(RNAi)* embryos, poleward chromosome scattering was so pronounced that the spindle equator became temporarily devoid of chromosomes; yet chromosomes eventually reversed direction and resumed congression (Fig. 2A, B; Movie S4). We conclude that PP4 inhibition produces a defect at the time of NEBD that precludes rapid chromosome bi-orientation and instead leads to an extended mono-oriented state. The defect is not permanent, however, because significant bi-orientation is eventually achieved later in prometaphase, albeit at the cost of attachment errors that compromise the fidelity of chromosome segregation.

**Figure 2.**
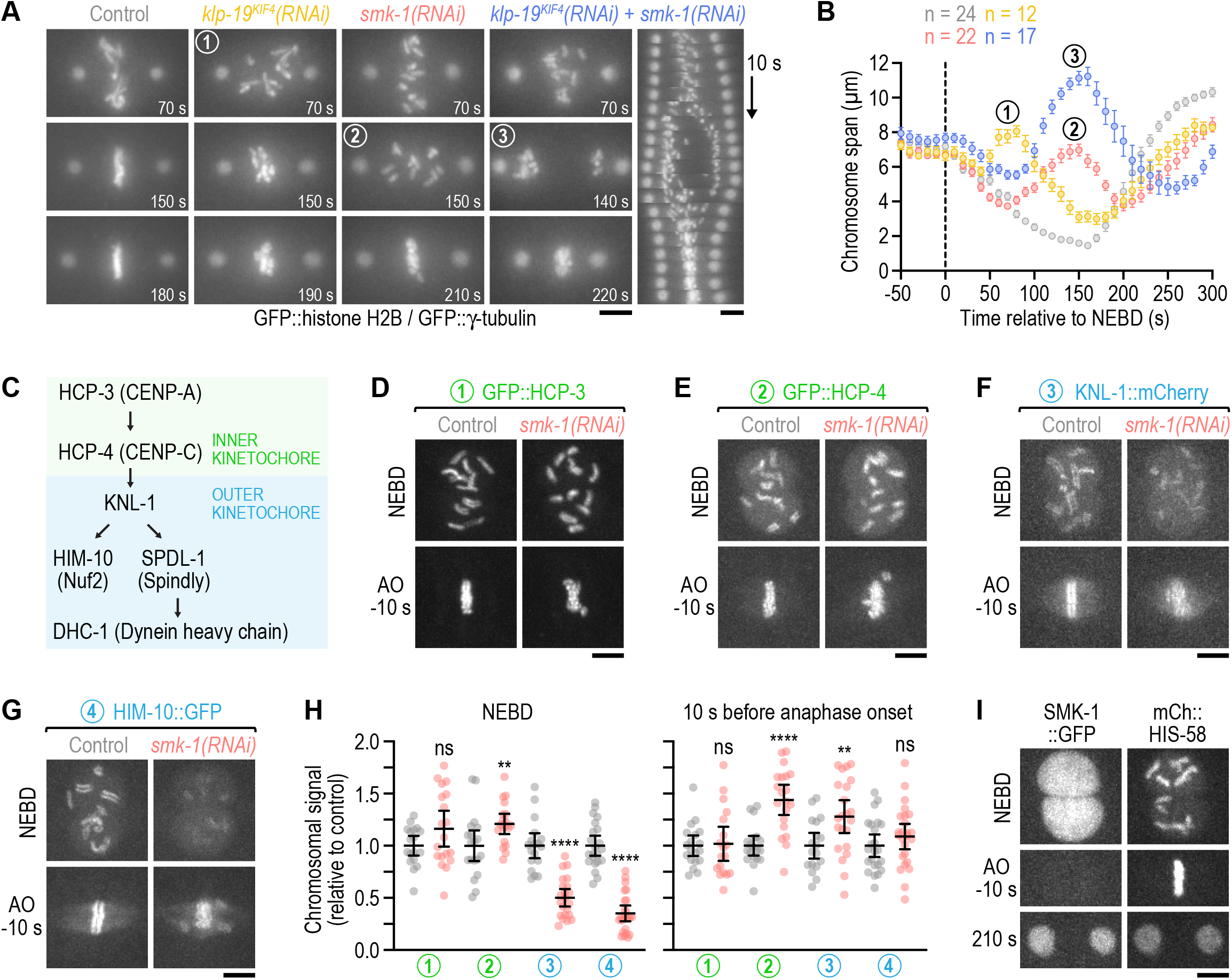
PP4 inhibition diminishes outer but not inner kinetochore assembly prior to NEBD. **(A)** Selected images and kymograph from time-lapse movies of one-cell embryos co-expressing GFP::histone H2B (HIS-58) and GFP::γ-tubulin (TBG-1). Time is relative to NEBD. Scale bar, 5 µm. **(B)** Chromosome span versus time relative to NEBD (mean of *n* embryos ± SEM). Measurements were performed in time-lapse movies such as those shown in *(A)*. Numbers in *(A)* and *(B)* refer to corresponding time points in images and graph. **(C)** Assembly hierarchy of the kinetochore components analyzed in this figure and *Fig. S2*. **(D) - (G)** Selected images from time-lapse movies of one-cell embryos co-expressing fluorescently tagged kinetochore components (GFP::HCP-3, GFP::HCP-4, and HIM-10::GFP are endogenously tagged; KNL-1::mCherry is a functional transgene) and transgenic GFP- or mCherry-tagged histone H2B (HIS-58). Only the kinetochore component is shown. Time points correspond to NEBD and the last frame prior to anaphase onset (AO). Scale bars, 5 µm. **(H)** Intensity of the chromosomal signal at NEBD or just prior to AO in the one-cell embryo (mean ± 95 % CI), normalized to the mean of the respective control, for the components shown in *(D)* - *(G)*. Statistical significance was determined by the Mann-Whitney test. \*\*\*\**P* < 0.0001; \*\**P* < 0.01; *ns* = not significant, *P* > 0.05. **(I)** Selected images from a time-lapse movie of a one-cell embryo co-expressing endogenously tagged SMK-1::GFP and transgenic mCherry::histone H2B (HIS-58). Scale bar, 5 µm.

### PP4 inhibition diminishes outer but not inner kinetochore assembly prior to NEBD

We next set out to understand why PP4 inhibition precludes rapid bi-orientation after NEBD. Live imaging in one-cell embryos showed that PP4 inhibition had no discernable effect on centrosome maturation (Fig. 1B, C, E, G; Fig. S1F, I), which contrasts with the conclusion of a prior study (Sumiyoshi *et al*., 2002). Consistent with this, PP4 inhibition affected neither pronuclear migration nor assembly and positioning of the mitotic spindle. The chromosome congression and segregation defects observed following PP4 inhibition are therefore not attributable to mis-regulation of the microtubule cytoskeleton.

We noticed that PP4 inhibition in KLP-19^Kif4^-depleted embryos suppressed the kinetochore-powered poleward chromosome movements in early prometaphase that are typically observed after KLP-19^Kif4^ loss (Powers *et al*., 2004; Fig. 2A, B; Movie S4). This suggested that PP4 inhibition prevented kinetochores from promptly engaging spindle microtubules at NEBD. We therefore examined the effect of PP4 inhibition on kinetochore assembly (Fig. 2C). Kinetochores are built in a hierarchical manner starting with the inner kinetochore components HCP-3^CENP-A^ and HCP-4^CENP-C^, which through HCP-4^CENP-C^ recruit components of the outer kinetochore, including the scaffolding protein KNL-1, the Ndc80 complex (represented here by the subunit HIM-10^Nuf2^), and the dynein module (represented here by SPDL-1^Spindly^ and dynein heavy chain DHC-1) (Cheeseman *et al*., 2004; Desai *et al*., 2003; Gassmann *et al*., 2008; Oegema *et al*., 2001). Using functional fluorescent versions of these components (Barbosa *et al*., 2017; Cheerambathur *et al*., 2019; Espeut *et al*., 2012; Yamamoto *et al*., 2008) we confirmed that control embryos assemble a robust outer kinetochore prior to NEBD (Fig. 2D - G). In *smk-1(RNAi)* embryos, chromosomal levels of GFP::HCP-3^CENP-A^ and GFP::HCP-4^CENP-C^ were similar and higher at NEBD than in controls, respectively (Fig. 2D, E, H; Movie S5). By contrast, chromosomal levels of outer kinetochore components were significantly lower at NEBD than in controls but subsequently recovered after NEBD, approximating control levels by anaphase onset (Fig. 2F - H; Movie S6). Thus, PP4 inhibition diminishes outer kinetochore assembly downstream of GFP::HCP-4^CENP-C^ and does so specifically prior to NEBD. Importantly, non-chromosomal levels of outer kinetochore components in prophase nuclei were increased in *smk-1(RNAi)* relative to controls (Fig. S2A), ruling out defective nuclear import as the cause for diminished outer kinetochore assembly. Chromosomal levels of SPDL-1^Spindly^ and dynein, which assemble on outer kinetochores after NEBD, eventually surpassed control levels in *smk-1(RNAi)* embryos (Fig. S2B - D; Movie S7), underscoring that PP4 inhibition delays outer kinetochore assembly rather than prevents it.

The effect of *smk-1(RNAi)* on prophase kinetochore assembly implies that PP4 acts in the nucleus. In agreement with this, live imaging of one-cell embryos expressing endogenously tagged SMK-1::GFP revealed that SMK-1 is strongly enriched in the male and female pronucleus (Fig. 2I; Movie S8). The nuclear signal remained diffuse throughout prophase with no detectable enrichment on condensing chromosomes. After NEBD, SMK-1::GFP gradually dissipated from the spindle region, equilibrated with the cytoplasmic signal by metaphase, and rapidly concentrated in nascent daughter nuclei at mitotic exit. We also added tags to PPH-4.1, but homozygous animals were either very sick (C-terminal 3xFLAG or GFP) or inviable (N-terminal 3xFLAG), precluding conclusive localization analysis.

Taken together, these results suggest that nuclear-enriched PP4 acts on prophase chromosomes to ensure timely outer kinetochore assembly prior to NEBD. In PP4-inhibited embryos, suppression of outer kinetochore assembly prior to NEBD, and subsequent relief of this suppression after nuclear dissolution, presumably by a cytoplasmic phosphatase activity, likely accounts for the unusual chromosome scattering and recovery phenotype.

### Delayed outer kinetochore assembly following PP4 inhibition also delays sister centromere resolution

Our analysis of kinetochore assembly showed that PP4 inhibition affects prophase kinetochore assembly downstream of HCP-4^CENP-C^. Since HCP-4^CENP-C^ is not only required for kinetochore assembly but also for resolution of sister centromeres (Moore and Roth, 2001; Moore *et al*., 2005), we analyzed sister centromere resolution following PP4 inhibition. We monitored sister centromere resolution using GFP::HCP-3^CENP-A^, which localizes to centromeres upstream of HCP-4^CENP-C^, and developed a live imaging assay to quantify the extent of resolution. This revealed that sister centromeres in PP4-inhibited embryos are poorly resolved when examined within the first minute after NEBD but are fully resolved by the time chromosomes have scattered later in prometaphase (Fig. 3A - D; Fig. S3A; Movie S9). Importantly, mitotic chromosomes in PP4-inhibited embryos had a compact morphology and were in fact slightly shorter than in controls (Fig. S3B, C), demonstrating that the resolution defect is not a byproduct of defective chromosome condensation. Moreover, in contrast to *wapl-1(RNAi)*, PP4 inhibition did not increase chromosomal levels of the cohesin subunit HIM-1^SMC1^::GFP (Fig. S3D, E), suggesting that the resolution defect is not caused by impaired cohesin removal. Instead, KNL-1 depletion revealed that outer kinetochore assembly is a prerequisite for sister centromere resolution (Fig. 3E, F; Fig. S3A; Movie S10). We therefore conclude that failure to assemble an outer kinetochore contributes to the sister centromere resolution defect that was previously reported for HCP-4^CENP-C^ depletion and that we confirm in this study (Fig. 3F; Fig. S3A). Taken together, these results suggest that the sister centromere resolution defect in PP4-inhibited embryos is a consequence of diminished outer kinetochore assembly during prophase and that recovery of outer kinetochore assembly after NEBD also restores sister centromere resolution.

**Figure 3.**
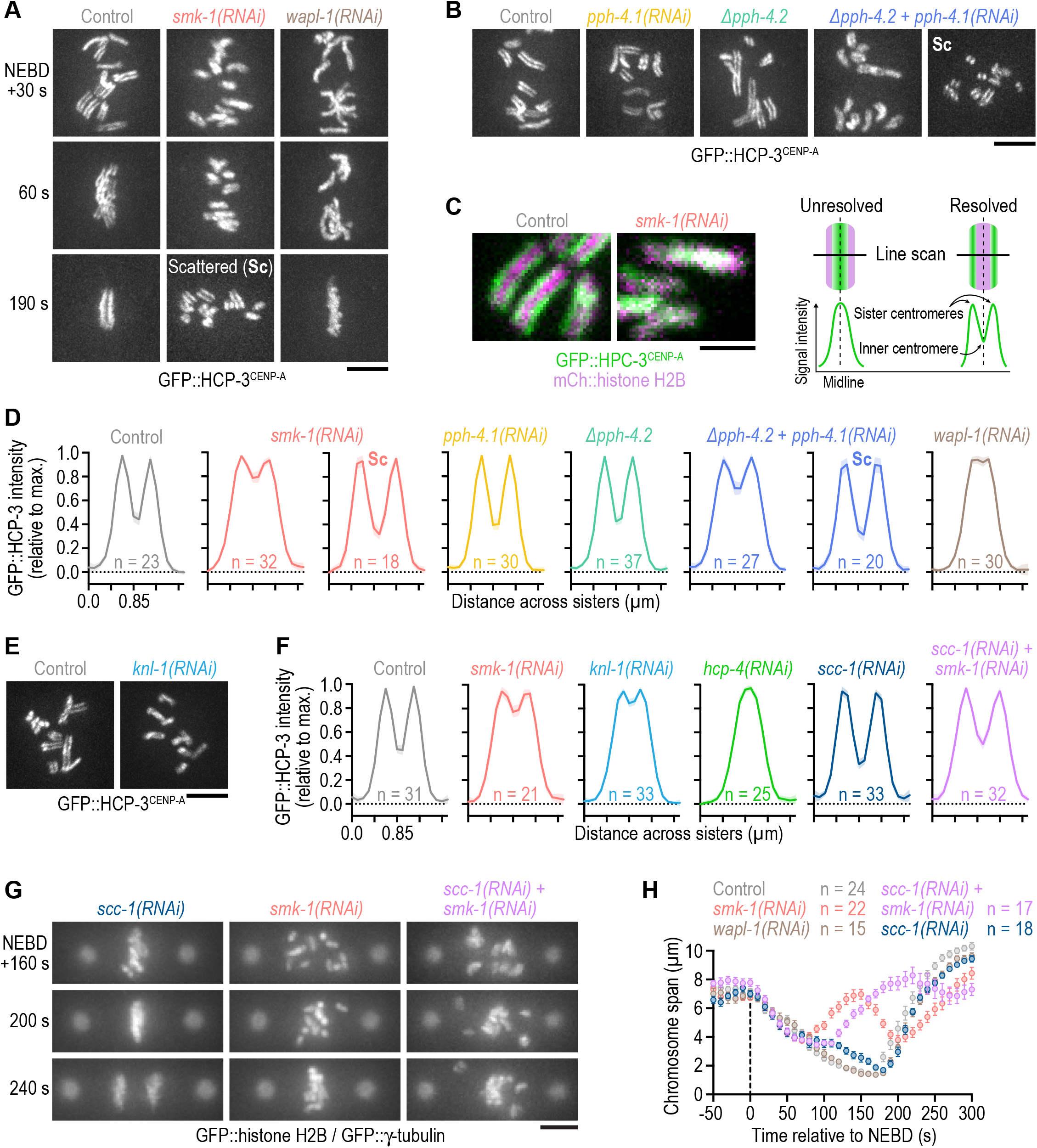
Delayed outer kinetochore assembly following PP4 inhibition also delays sister centromere resolution. **(A), (B)** Selected images from time-lapse movies of one-cell embryos expressing endogenously tagged GFP::CENP-A (HCP-3) to mark sister centromeres. Time in *(A)* is relative to NEBD. Time point in *(B)* corresponds to 30 s after NEBD. For *∆pph-4*.*2 + pph-4*.*1(RNAi)*, the time of chromosome scattering (Sc) is also shown. Scale bars, 5 µm. **(C)** *(left)* Selected images from time-lapse movies of one-cell embryos co-expressing endogenously tagged GFP::HCP-3 and transgenic mCherry::histone H2B (HIS-58). Time is relative to NEBD. Scale bar, 2 µm. *(right)* Schematic illustrating how line scans of the GFP::HCP-3 signal across the short axis of condensed mitotic chromosomes report on the extent of sister centromere resolution. **(D), (F)** Line scan profiles (mean of *n* chromosomes from at least 10 embryos ± 95 % CI), generated as described in *(C)*. Profiles in *(D)* are of endogenously tagged GFP::HCP-3, and profiles in *(F)* are of transgenic GFP::HCP-3 in an *hcp-3* knock-out background. **(E)** Selected images from time-lapse movies of one-cell embryos expressing transgenic GFP::HCP-3. Time point corresponds to 30 s after NEBD. Scale bar, 5 µm. **(G)** Selected images from time-lapse movies of one-cell embryos co-expressing GFP::histone H2B (HIS-58) and GFP::γ-tubulin (TBG-1). Time is relative to NEBD. Scale bar, 5 µm. **(H)** Chromosome span (mean of *n* embryos ± SEM) versus time relative to NEBD. Measurements were performed in embryos such as those shown in *(G)*.

To address whether delayed sister centromere resolution contributes to poleward chromosome scattering in PP4-inhibited embryos, we asked whether inhibiting sister centromere resolution by preventing cohesin removal also results in chromosome scattering, and whether rescuing prophase sister centromere resolution in PP4-inhibited embryos by depleting cohesin prevents chromosome scattering. Despite the failure to resolve sister centromeres due to defective cohesin removal (Fig. 3A, D; Fig. S3A, D, E), chromosomes in *wapl-1(RNAi)* embryos congressed normally and formed a tight metaphase plate (Fig. 3A, H). Thus, chromosomes can congress in a timely manner even when sister centromere resolution is substantially impaired. Moreover, depletion of the cohesin subunit SCC-1 rescued sister centromere resolution following PP4 inhibition but failed to rescue chromosome congression (Fig. 3F - H; Fig. S3A). Taken together, these results argue against the idea that delayed sister centromere resolution is causative for the chromosome congression defects in PP4-inhibited embryos. However, delayed sister centromere resolution likely contributes to chromosome mis-segregation in PP4-inhibited embryos by elevating merotelic attachments, in which a kinetochore is simultaneously connected to both spindle poles.

### HCP-4^CENP-C^ binding to SMK-1 is dispensable for chromosome segregation

Our results show that PP4 inhibition delays outer kinetochore assembly and sister centromere resolution, both of which require HCP-4^CENP-C^. Prior work in *D. melanogaster* has shown that CENP-C binds the SMK-1 homolog Falafel and becomes hyperphosphorylated following PP4 inhibition (Lipinszki *et al*., 2015). Thus, an attractive hypothesis is that HCP-4^CENP-C^ recruits PP4 via an interaction with SMK-1 and is a potential PP4 substrate, and that the outer kinetochore assembly defect in PP4-inhibited embryos reflects defective PP4 targeting to HCP-4^CENP-C^. To test this idea, we first asked whether HCP-4^CENP-C^ binds SMK-1. Like Falafel, SMK-1 contains an EVH1 domain near its N-terminus (Fig. 4A). Using purified recombinant proteins, we performed GST pull-downs with full-length HCP-4^CENP-C^ and the GST-tagged EVH1 domain (residues 146-282). Full-length HCP-4^CENP-C^ bound to GST::EVH1, and mutating the conserved EVH1 residues Y179 and W187 to alanine abrogated the interaction (Fig. 4B, C). Thus, the EVH1 domain of SMK-1 binds FxxP motifs in the same manner as the EVH1 domains of the fly and human homologs (Lipinszki *et al*., 2015; Ueki *et al*., 2019).

**Figure 4.**
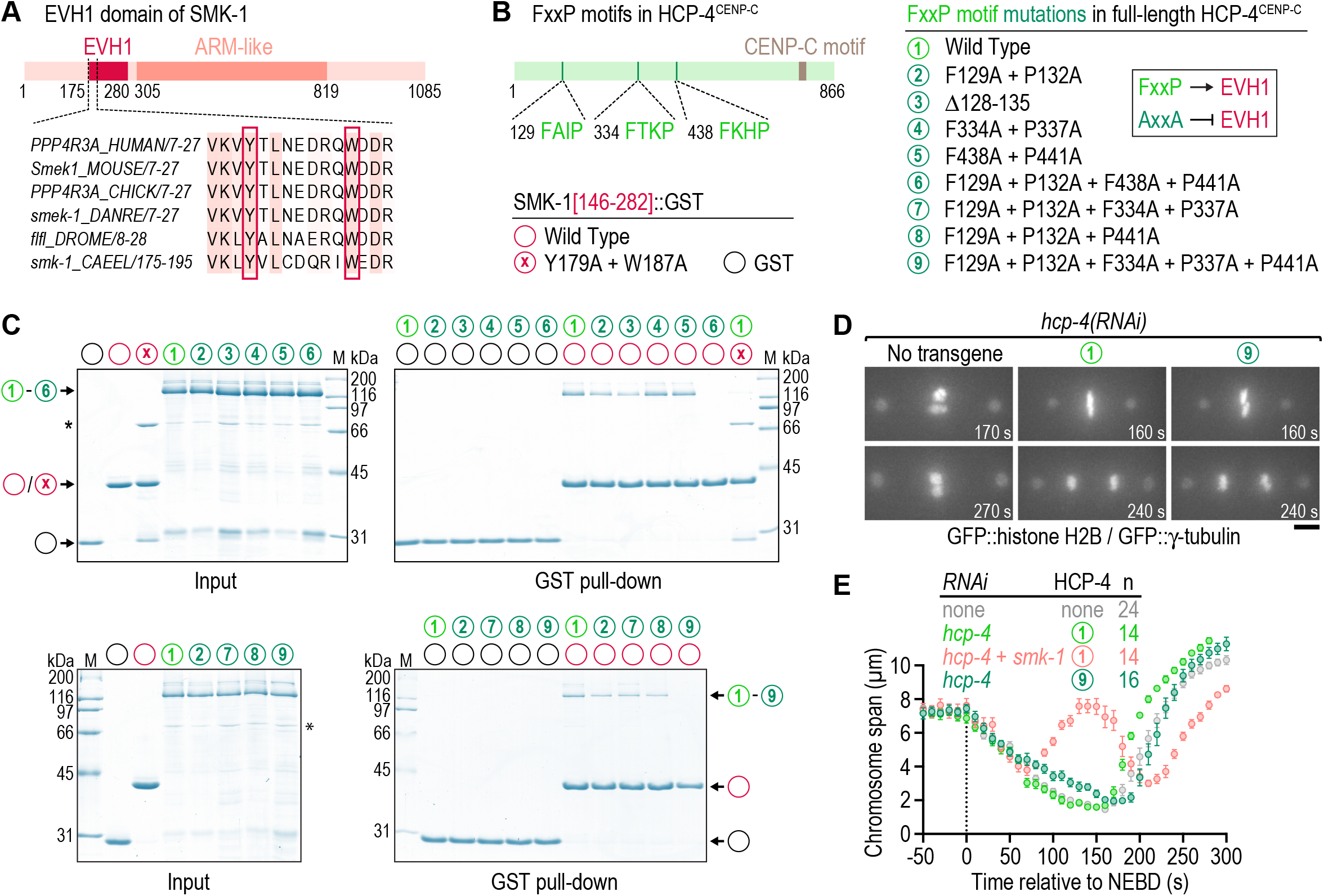
HCP-4^CENP-C^ binding to SMK-1 is dispensable for chromosome segregation. **(A)** Schematic of SMK-1 and sequence alignment in the highly conserved EVH1 domain. The tyrosine and tryptophan residues required for FxxP motif binding are highlighted. **(B)** Schematic of HCP-4^CENP-C^ showing the locations of FxxP motifs and overview of HCP-4 and SMK-1 mutations evaluated in pull-down assays. Mutating FxxP motifs to AxxA prevents their interaction with the EVH1 domain, as indicated in the cartoon. **(C)** Coomassie Blue-stained protein gels showing purified recombinant proteins (Input) and proteins eluted from glutathione agarose resin after GST pull-down. Molecular weight of size markers (M) is indicated in kilodaltons (kDa). **(D)** Selected images from time-lapse movies of one-cell embryos co-expressing GFP::histone H2B (HIS-58) and GFP::γ-tubulin (TBG-1). Time is relative to NEBD. Numbers refer to transgene-encoded wild-type and mutant HCP-4, as described in *(B)*. Scale bar, 5 µm. **(E)** Chromosome span (mean of *n* embryos ± SEM) versus time relative to NEBD. Measurements were performed in time-lapse movies such as those shown in *(D)*.

HCP-4^CENP-C^ contains three FxxP motifs in its N-terminal half (Fig. 4B), an arrangement that contrasts with the single C-terminally located Falafel-interacting FxxP motif of *D. melanogaster* CENP-C. HCP-4^CENP-C^ in which any of the three FxxP motifs was mutated to AxxA (or in which the first FxxP motif was deleted) still bound GST::EVH1, as did HCP-4^CENP-C^ in which the first and second FxxP motifs were mutated to AxxA (Fig. 4B, C). By contrast, the combination of first and third AxxA abrogated the interaction (mutant number 6; Fig. 4B), as did the combination of first AxxA, second AxxA, and third FxxA (mutant number 9). Furthermore, *in vitro* translated HCP-4^CENP-C^ fragments containing individual FxxP motifs bound to GST::EVH1 (data not shown). We conclude from these binding experiments that all three FxxP motifs in HCP-4^CENP-C^ participate in the interaction with SMK-1.

To test whether SMK-1 binding to HCP-4^CENP-C^ is important for outer kinetochore assembly prior to NEBD, we set up a molecular replacement system based on transgenic expression of RNAi-resistant mCherry::HCP-4^CENP-C^ from a defined chromosomal locus after single-copy integration (Fig. S3F). Depletion of endogenous HCP-4^CENP-C^ resulted in complete failure to segregate chromosomes, as expected, and was rescued by RNAi-resistant wild-type mCherry::HCP-4^CENP-C^ (Fig. 4D). For the SMK-1 binding-defective HCP-4^CENP-C^ mutant we chose mutant number 9 rather than number 6 (Fig. 4B), because mutating the phenylalanine in the third FxxP motif to alanine unexpectedly interfered with nuclear localization of mCherry::HCP-4^CENP-C^ (data not shown). The SMK-1 binding-defective HCP-4^CENP-C^ mutant supported near-normal chromosome congression, timely sister centromere resolution, and error-free chromosome segregation (Fig. 4D, E; Fig. S3G), implying that the PP4 interaction with HCP-4^CENP-C^ mediated by the EVH1 domain of SMK-1 is dispensable for normal mitosis. These data suggest either that HCP-4^CENP-C^ is not a relevant PP4 target or that PP4 is sufficiently concentrated in the nucleus for HCP-4^CENP-C^ dephosphorylation to occur in the absence of the SMK-1–HCP-4^CENP-C^ interaction.

### The Ndc80-Ska module is essential for recovery from mono-orientation induced by kinetochore dynein following PP4 inhibition

Scattered chromosomes in PP4-inhibited one-cell embryos eventually resume congression, even when initial mono-orientation is exacerbated by co-depletion of KLP-19^Kif4^ (Fig. 2A, B; Movie S4), and form bi-oriented attachments capable of sustaining tension (a fraction of these being merotelic). We therefore sought to address the mechanism responsible for this striking recovery. The microtubule-binding Ska complex (a trimer that in *C. elegans* consists of two SKA-1 subunits and one SKA-3 subunit) has emerged as a conserved attachment maturation factor at kinetochores (Cheerambathur *et al*., 2017; Hanisch *et al*., 2006; Janczyk *et al*., 2017; Schmidt *et al*., 2012; Welburn *et al*., 2009). To test whether Ska promotes bi-orientation of scattered chromosomes, we depleted SMK-1 in *∆ska-1* embryos. Loss of SKA-1 on its own caused a very mild chromosome congression defect. By contrast, *∆ska-1;smk-1(RNAi)* completely blocked re-congression of scattered chromosomes, which remained close to spindle poles until the end of mitosis (Fig. 5A, B; Movie S11). Consistent with a persistent mono-oriented state of *∆ska-1;smk-1(RNAi)* chromosomes, spindle poles in the co-inhibition separated prematurely without a subsequent recovery of spindle length. The same phenotype was observed in *ska-3(RNAi);smk-1(RNAi)* embryos, and examination of GFP::HCP-3^CENP-A^ confirmed that sister chromatids co-segregate to the same spindle pole prior to their separation (Fig. 5C). Consistent with Ska being important for recovery from initial mono-orientation, enhancing Ska recruitment to kinetochores by introducing phosphorylation site mutations in NDC-80 (Cheerambathur *et al*., 2017) ameliorated chromosome scattering following PP4 inhibition (Fig. 5D).

**Figure 5.**
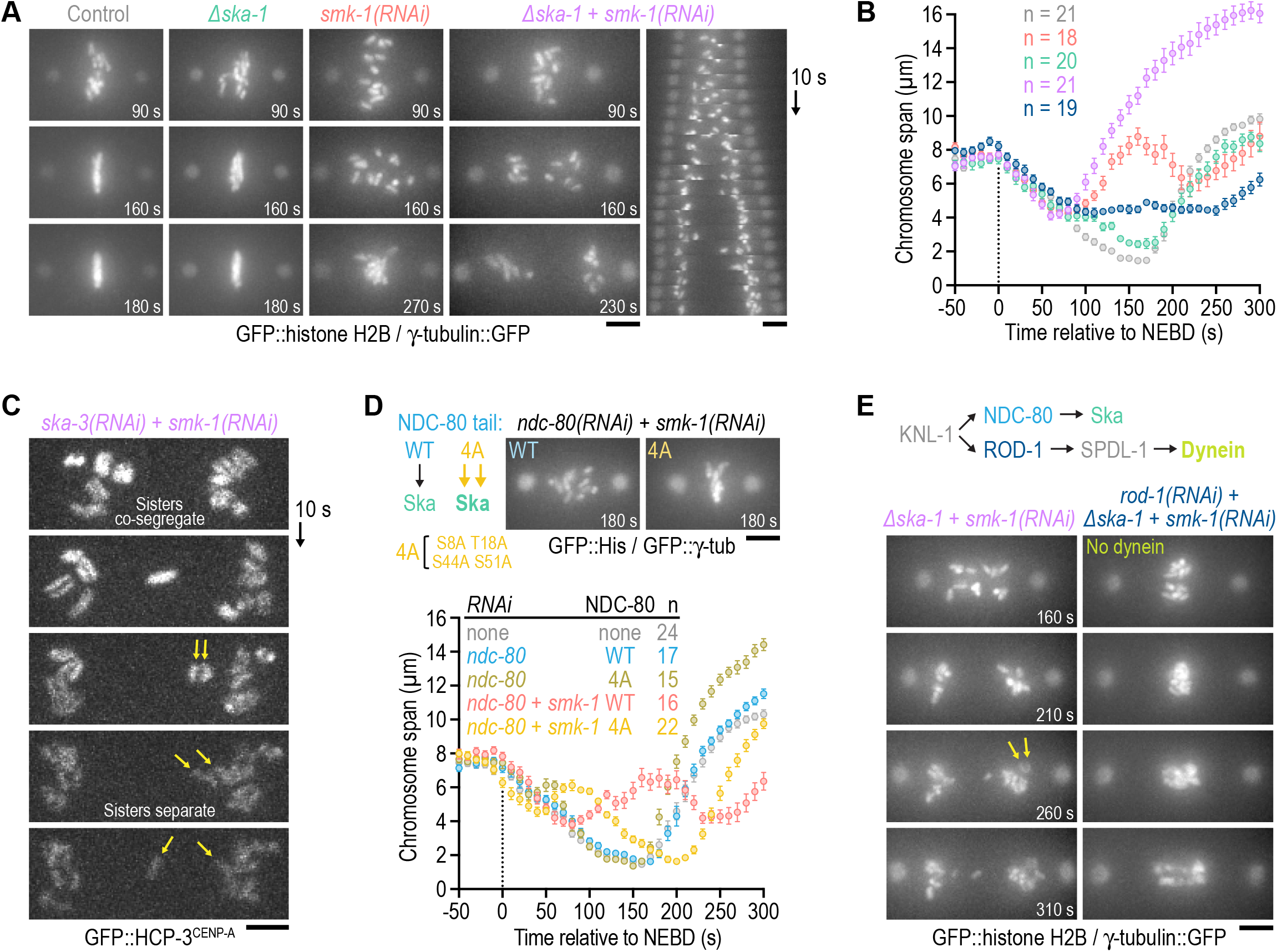
The Ndc80-Ska module is essential for recovery from mono-orientation induced by kinetochore dynein following PP4 inhibition. **(A), (E)** Selected images from time-lapse movies of one-cell embryos co-expressing GFP::histone H2B (HIS-11) and γ-tubulin::GFP (TBG-1). Time is relative to NEBD. A kymograph for *∆ska-1 + smk-1(RNAi)* is also shown in *(A)*. Cartoon in *(E)* shows recruitment hierarchy at the outer kinetochore relevant for this figure. The Arrows in *(E)* point to separating sister chromatids. Scale bars, 5 µm. **(B)** Chromosome span (mean of *n* embryos ± SEM) versus time relative to NEBD. Measurements were performed in embryos such as those shown in *(A)*. Conditions are color-coded as in *(A)* and *(E)*. **(C)** Successive frames from a time-lapse movie of a one-cell embryo expressing endogenous GFP::HCP-3 to mark centromeres. Top frame corresponds to late prometaphase. Arrows point to a pair of co-segregated sister chromatids which later separate. Scale bar, 2 µm. **(D)** *(top left)* Cartoon illustrating that phosphorylation site mutations in NDC-80’s N-terminal tail promote Ska recruitment to kinetochores. *(top right)* Selected images from time-lapse movies of one-cell embryos expressing transgene-encoded wild-type (WT) or mutant (4A) NDC-80 in the background of GFP::histone H2B (HIS-58) and GFP::γ-tubulin (TBG-1). Time is relative to NEBD. Scale bar, 5 µm. *(bottom)* Chromosome span (mean of *n* embryos ± SEM) versus time relative to NEBD, measured in embryos such as those shown above this graph.

We next investigated the mechanism leading to mono-orientation of chromosomes in PP4-inhibited embryos that persists when Ska is inhibited. A potential candidate driver of this mono-orientation is kinetochore dynein, whose recruitment requires the RZZ complex (comprised of ROD-1, ZWL-1^Zwilch^ and CZW-1^Zw10^) and SPDL-1^Spindly^ (Gassmann *et al*., 2008; Fig. 5E). Consistent with this idea, the persistent mono-orientation and sister chromatid co-segregation in *∆ska-1;smk-1(RNAi)* embryos was suppressed by depleting ROD-1 to prevent kinetochore dynein recruitment: chromosomes in *∆ska-1;smk-1(RNAi)*;*rod-1(RNAi)* embryos remained near the spindle equator and became stretched out along the spindle axis in anaphase (Fig. 5E; Movie S12). Thus, kinetochore dynein is required for chromosome scattering following PP4 inhibition. This likely reflects dynein’s ability to power poleward chromosome movement and/or promote mono-orientation by limiting the formation of bipolar merotelic attachments (Gassmann *et al*., 2008; Edwards *et al*., 2018). The prominent stretching of chromosomes in anaphase indeed suggests that most kinetochores are merotelically attached in *∆ska-1;smk-1(RNAi)*;*rod-1(RNAi)* embryos (Fig. 5E).

Collectively, these data establish that the outer kinetochore assembly defect prior to NEBD in PP4-inhibited embryos results in an imbalance of kinetochore-based microtubule-interfacing activities after NEBD, with kinetochore dynein being responsible for aberrant mono-orientation and poleward scattering followed by subsequent bi-orientation mediated by the Ndc80-Ska module. Notably, this imbalance results in extensive merotely and mis-segregation, highlighting the critical importance of properly timed outer kinetochore assembly for faithful mitosis.

## Conclusion

Here we show that nuclear-enriched PP4 is critical for properly timed outer kinetochore assembly and that this in turn is important for the proper order and fidelity of chromosome-microtubule interactions. It is interesting that a phosphatase is important for this function, as mitotic assembly reactions are typically promoted by phosphorylation and reversed by dephosphorylation. While the regulatory EVH1 domain-containing subunit of PP4 complexes is known to interact with the inner kinetochore protein CENP-C, this interaction seems dispensable for timely outer kinetochore assembly. Thus, a major future goal will be to address the key substrate(s) of PP4 whose dephosphorylation underlies this critical event ensuring orderly and accurate execution of mitosis. Our analysis of PP4 has also uncovered the importance of outer kinetochore assembly for sister centromere resolution, a critical event in ensuring properly bioriented attachments. Thus, timely outer kinetochore assembly is also important to restrain the frequency of merotelic attachments that are responsible for chromosome mis-segregation. Finally, our results show that the Ndc80-Ska module can promote the conversion from mono- to bi-orientation. We propose that this function of Ndc80-Ska is ordinarily masked by mechanisms that ensure rapid bi-orientation in early prometaphase but are suppressed in PP4-inhibited embryos because outer kinetochore assembly is diminished in prophase. For example, polar ejection forces mediated by the chromokinesin KLP-19^Kif4^ are strongest at NEBD, i.e. when chromosomes are closest to spindle poles (Ke *et al*., 2009), and are proposed to promote bi-orientation by generating torque on mono-oriented chromosomes such that sister kinetochores come to face opposite poles (Powers *et al*., 2004). Although the Ndc80-Ska module can compensate for the failure of kinetochores to rapidly engage microtubules at NEBD by promoting bi-orientation later in prometaphase, overreliance on this pathway produces an unacceptable number of attachment errors and is incompatible with proper chromosome segregation. PP4 inhibition therefore demonstrates that outer kinetochore assembly prior to NEBD is crucial for error-free mitosis chromosome segregation.

## MATERIALS AND METHODS

### *Caenorhabditis elegans* strains

Worm strains (Table S1) were maintained at 20 ºC on standard nematode growth media (NGM) plates seeded with OP50 bacteria. A Mos1 transposon-based strategy (MosSCI; Frøkjær-Jensen *et al*., 2012) was used to generate strains stably expressing mCherry::HCP-4 under the control of the *mex-5* promoter and tbb-2 3′ UTR for expression in germline cells. Transgenes were cloned into pCFJ352 for insertion on Chromosome 1 (*ttTi4348* locus) or pCFJ151 for insertion on Chromosome 2 (*ttTi5605* locus), and transgene integration was confirmed by PCR and sequencing. Other fluorescent markers were subsequently introduced by mating.

The alleles *smk-1::3xflag, smk-1::gfp, ∆smk-1*, and *∆pph-4*.*2* were generated by CRISPR/Cas9-mediated genome editing, as described previously (Arribere *et al*., 2014; Paix *et al*., 2014). Genomic sequences targeted by sgRNAs are listed in Table S2. Screening was performed by PCR and edits were confirmed by sequencing. Strains were outcrossed 4 - 6 times against the wild-type N2 strain to remove potential background mutations, and fluorescent markers were subsequently introduced by mating. Homozygous *∆pph-4*.*2* animals are fully viable. By contrast, homozygous *∆smk-1* and *∆pph-4*.*1* animals (F1) grow to adulthood and produce embryos that are inviable (Fig. S1C, D). *∆smk-1* animals additionally become sterile soon after they start to produce embryos, which are very fragile and remain inside the mother. *∆smk-1* and *∆pph-4*.*1* were therefore maintained in a heterozygous state using a GFP-marked genetic balancer, and homozygous F1 progeny from heterozygous mothers were identified by the absence of GFP fluorescence in the pharynx.

### RNA interference

For production of double-stranded RNA (dsRNA), oligonucleotides with tails containing T3 and T7 promoters (Table S3) were used to amplify regions from wild-type N2 cDNA (*ndc-80* and *pph-4*.*1*) or genomic DNA (all other genes). PCR products were purified (NucleoSpin Gel and PCR Clean-up, Macherey-Nagel) and used as templates for T3 and T7 transcription reactions (MEGAscript, Invitrogen). Transcription reactions were purified (NucleoSpin RNA Clean-up, Macherey-Nagel) and annealed in 3x soaking buffer (32.7 mM Na_2_HPO_4_, 16.5 mM KH_2_PO_4_, 6.3 mM NaCl, 14.1 mM NH_4_Cl). dsRNAs were delivered by injecting L4 hermaphrodites. After injection, animals were kept on NGM plates seeded with OP50 bacteria for 48 h at 20 ºC before embryo progeny was isolated for analysis. For strain TH66 (Fig. S1I), injected animals were kept at 25 ºC for 24 - 30 h to prevent silencing of the *gfp::ebp-2* transgene. As a control for the *smk-1(RNAi);rod-1(RNAi)* double depletion (Fig. 5D), *smk-1(RNAi)* alone was performed with a 1:1 dilution of dsRNA targeting *smk-1* with dsRNA targeting *C09B9*.*4*, which does not have any obvious function in the one-cell embryo.

### Live imaging

Gravid hermaphrodites were dissected in a watch glass filled with a 0.7× dilution of Egg Salts medium (1× medium is 118 mM NaCl, 40 mM KCl, 3.4 mM MgCl_2_, 3.4 mM CaCl_2_, 5 mM HEPES pH 7.4). Embryos were mounted on a 2 % agarose pad and covered with an 18 mm × 18 mm coverslip (No. 1.5H, Marienfeld). All imaging was performed in temperature-controlled rooms kept at 20 ºC. Two microscopes were used: a Zeiss Axio Observer microscope, equipped with an Orca Flash 4.0 camera (Hamamatsu) and a Colibri 2 light source (Zeiss), controlled by ZEN software (Zeiss); and a Nikon Eclipse Ti microscope coupled to an Andor Revolution XD spinning disk confocal system, composed of an iXon Ultra 897 CCD camera (Andor Technology), a solid-state laser combiner (ALC-UVP 350i, Andor Technology), and a CSU-X1 confocal scanner (Yokogawa Electric Corporation), controlled by Andor IQ3 software (Andor Technology).

### Image acquisition and analysis

#### Chromosome span and pole-pole distance

Time-lapse movies of one-cell embryos co-expressing GFP::HIS-58/GFP::TBG-1 (Fig. 1D; Fig. 2B; Fig. 3H; Fig. 4E; Fig. 5D), GFP::HIS-11/TBG-1::GFP (Fig. 5B), or mCherry::HIS-11/GFP::TBB-2 (Fig. 1F; Fig. S1G) were acquired on the Axio Observer microscope with a 63x NA 1.4 Plan-Apochromat objective (Zeiss) at 2 × 2 binning. A 9 × 1.5 µm z-stack was captured every 10 s from just prior to NEBD until chromosome decondensation, and maximum intensity projections of z-stacks were used for measurements in Fiji (Image J version 2.0.0-rc-56/1.53 c).

For chromosome span and pole-pole distance plots, the x and y coordinate of centrosomes, and of the chromosomes closest to each centrosome, were recorded for each frame using the MTrackJ plugin by manually clicking in the center of centrosomes and on the outer edge of chromosomes. Chromosome span was defined as the distance between the two outermost chromosomes measured along the spindle axis. Tracks from individual embryos were aligned relative to NEBD, defined as the first frame with a decrease in the diffuse nuclear histone signal due to equilibration with the cytoplasmic pool. Anaphase onset was defined as the first frame with discernible sister chromatid separation. Replicate profiles were aligned relative to the time of NEBD and plotted as mean ± SEM.

#### Anaphase chromatin bridges

Time-lapse movies of one-cell embryos co-expressing GFP::HIS-58/GFP::TBG-1 (Fig. 1B) or mCherry::HIS-11/GFP::TBB-2 (Fig. 1E; Fig. S1F) were acquired on the spinning disk confocal microscope with a 60x NA 1.4 Plan-Apochromat objective (Nikon) at 1 × 1 binning. A 9 × 1 µm z-stack was captured every 10 s from just prior to NEBD until chromosome decondensation, and maximum intensity projections were analyzed in Fiji. Anaphases were scored as having chromatin bridges if GFP::HIS-58 or mCherry::HIS-11 signal connecting the two masses of segregating chromosomes was visible three frames after the onset of sister chromatid separation.

#### Chromosomal and nuclear levels

Time-lapse movies of one-cell embryos expressing GFP::HCP-3/mCherry::HIS-58, GFP::HCP-4, KNL-1::mCherry/GFP::HIS-58, HIM-10::GFP/mCherry::HIS-58, GFP::SPDL-1/mCherry::HIS-58, or HIM-1::GFP/mCherry::HIS-58 were acquired on the spinning disk confocal microscope with a 60x NA 1.4 Plan-Apochromat objective (Nikon) at 1 × 1 binning. A 9 × 1 µm z-stack was captured every 10 s from just prior to NEBD until chromosome decondensation, and maximum intensity projections were analyzed in Fiji.

For analysis at NEBD (Fig. 2H *left*; Fig. S2A), the integrated intensity was measured within a region corresponding the area occupied by the nucleus, as well as within three smaller intranuclear regions that cumulatively covered a large fraction of the nuclear area not occupied by chromosomes (nuclear background). The chromosomal signal was calculated by subtracting the integrated nuclear background intensity (multiplied by the ratio of nuclear region area to background region area) from the integrated intensity of the nuclear region. The diffuse non-chromosomal nuclear signal (Fig. S2A) was calculated by subtracting the integrated background intensity measured in cytoplasmic regions adjacent to the nucleus from the integrated intensity of the three intranuclear regions not occupied by chromosomes. Signal intensities of replicates were averaged, normalized to the average of the respective control, and plotted as mean ± 95 % CI.

For analysis just prior to anaphase onset (Fig. 2H *right*; Fig. S3E), defined as one to three frames before sister chromatid separation, the integrated intensity was determined in the region encompassing the chromosomes. The chromosomal region was expanded by 2 pixels, and the difference in integrated intensity between the expanded and chromosomal region was used to define the background intensity. The chromosomal signal was calculated by subtracting the integrated background intensity (multiplied by the ratio of chromosomal region area to background region area) from the integrated intensity of the chromosomal region. Signal intensities of replicates were averaged, normalized to the average of the respective control, and plotted as mean ± 95 % CI.

For analysis of chromosomal signal over time (Fig. S2C) the ‘Find Maxima’ function in Fiji was used to identify the top 15 local maxima in the region containing the chromosomes, and the values were averaged. The averaged intensity of the top 15 local maxima of an adjacent cytoplasmic region was subtracted as background. Signal intensities of replicate time-lapse series were averaged, normalized to the peak signal within the time-lapse series of the respective control, aligned relative to the time of NEBD, and plotted as mean ± SEM.

#### Sister centromere resolution

Time-lapse movies of one-cell embryos expressing endogenous GFP::HCP-3 (Fig. 3D; Fig. S3A *left*, G) or transgenic GFP::HCP-3 in an *hcp-3* knock-out background (Fig. 3F; Fig. S3A *right*) were acquired on the spinning disk confocal microscope with a 100x NA 1.45 Plan-Apochromat objective (Nikon) at 1 × 1 binning. A 9 × 1 µm z-stack was captured every 10 s from just prior to NEBD until chromosome decondensation, and maximum intensity projections were analyzed in Fiji. Measurements were made in the first 6 frames following NEBD, and, in PP4 inhibitions, also later in prometaphase when chromosomes had scattered.

For line scan plots (Fig. 3D, F; Fig. S3G), a 2-µm long and 3 pixel-wide line, drawn perpendicularly to the long axis of an individual chromosome, was centered on the chromosome midline and the fluorescence profile was determined using the ‘Plot Profile’ function in Fiji. Signal intensities were normalized to the maximum intensity within each profile. Profiles of replicates were averaged and plotted as mean ± 95 % CI.

For signal ratio plots (Fig. S3A, G), the intensity at the chromosome midline was divided by the maximum intensity within the line scan. Ratios of replicates were averaged, normalized to the average ratio of the respective control, and plotted as mean ± 95 % CI.

#### Chromosome size

Z-stacks with step size 0.1 µm were acquired in one-cell embryos co-expressing GFP::HIS-58/GFP::TBG-1 on the spinning disk confocal microscope with a 100x NA 1.45 Plan-Apochromat objective (Nikon) at 1 × 1 binning in late prophase or early prometaphase. Z-stacks were visualized in 3D using Imaris 9.5.0 (Bitplane). Chromosome length and width measurements (Fig. S3C) were made by visually delimiting chromosomes and assigning points at the extremities and in the middle of individual chromosomes. Replicate measurements were averaged and plotted as mean ± 95 % CI.

#### Microtubule nucleation

Time-lapse movies of one-cell embryos expressing EBP-2::GFP were acquired on the spinning disk confocal microscope with a 100x NA 1.45 Plan-Apochromat objective (Nikon) at 1 × 1 binning. A single central z-section that included both centrosomes was captured every 200 ms at metaphase for a total of 1 min.

For analysis of microtubule nucleation rates (Fig. S1I), an arc positioned 30 pixels away from the centrosome was drawn using the segmented line tool in Fiji, and a kymograph of at least 200 frames was generated using the Fiji Multi kymograph plugin. The number EBP-2::GFP puncta on the kymograph were manually counted and divided by the kymograph time axis. Replicate measurements were averaged and plotted as mean ± 95 % CI.

### Embryonic viability and brood size

Brood size (Fig. S1C) and embryonic viability assays (Fig. S1D) were performed at 20 ºC. Wild-type N2 L4 hermaphrodites, injected with dsRNA, or untreated L4 hermaphrodites of homozygous mutants, were grown for 40 h on NGM plates containing OP50 bacteria. Single adults were then placed on new plates with a small amount of OP50, removed 8 h later, and the plates were incubated for another 16 - 24 h to give viable embryos time to hatch. Embryonic viability was calculated by dividing the number of hatched larvae by the total number of progeny on the plate. Replicate counts were averaged and plotted as mean percentage ± 95 % CI.

### Plasmids for recombinant protein expression

cDNA encoding for full-length HCP-4 (UniProt entry G5EDA3) was inserted into a 2CT vector [N-terminal 6xHis::maltose binding protein (MBP) followed by a TEV protease cleavage site and C-terminal StrepTagII], and cDNA encoding for SMK-1 (UniProt entry H2KYN6; residues 146-282) was inserted into pGEX-6P-1 [N-terminal glutathione S-transferase (GST) followed by a Prescission protease cleavage site] containing C-terminal 6xHis.

### Recombinant protein expression and purification

Expression vectors were transformed into *E. coli* strains BL21 or Rosetta. Expression was induced at an OD_600_ of 0.9 with 0.1 mM IPTG. After expression overnight at 18 ºC, cells were harvested by centrifugation at 4000 × g for 20 min.

For purification of 6xHis::MBP::HCP-4::StrepTagII, bacterial pellets were resuspended in lysis buffer A (50 mM HEPES, 250 mM NaCl, 10 mM imidazole, 1 mM DTT, 1 mM PMSF, 2 mM benzamidine-HCl, pH 8.0), lysed by sonication, and cleared by centrifugation at 40000 × g for 45 min at 4 ºC. Proteins were purified by tandem affinity chromatography using HisPur Ni-NTA resin (Thermo Fisher Scientific) followed by strep-tactin sepharose resin (IBA). Ni-NTA resin was incubated in batch with the cleared lysate in the presence of 0.1 % (v/v) Tween 20 for 1 h at 4 ºC and washed with wash buffer A (25 mM HEPES, 250 mM NaCl, 25 mM imidazole, 0.1 % Tween 20, 1 mM DTT, 2 mM benzamidine-HCl, pH 8.0). Proteins were eluted on a gravity column with elution buffer A (50 mM HEPES, 150 mM NaCl, 250 mM imidazole, 1 mM DTT, 2 mM benzamidine-HCl, pH 8.0). Fractions containing the recombinant protein were pooled, incubated in batch with strep-tactin sepharose resin for 1 h at 4 ºC, and washed with wash buffer B (25 mM HEPES, 250 mM NaCl, 0.1 % Tween 20, 1 mM DTT, 2 mM benzamidine-HCl, pH 8.0). Proteins were eluted on a gravity column with elution buffer B (100 mM Tris-HCl, 150 mM NaCl, 1 mM EDTA, 10 mM desthiobiotin, pH 8.0), and the eluate was dialyzed against storage buffer (25 mM HEPES, 150 mM NaCl, pH 7.5). Glycerol and DTT were added to a final concentration of 10 % (v/v) and 1 mM, respectively, and aliquots were flash frozen in liquid nitrogen and stored at -80 ºC.

For purification of GST::SMK-1(146-282)::6xHis, bacterial pellets were resuspended in lysis buffer B (50 mM HEPES, 250 mM NaCl, 10 mM EDTA, 10 mM EGTA, 1 mM DTT, 1 mM PMSF, 2 mM benzamidine-HCl, pH 8.0), lysed by sonication, and cleared by centrifugation at 40000 × g for 45 min at 4 ºC. Proteins were purified by tandem affinity chromatography using glutathione agarose resin (Thermo Fisher Scientific) followed by Ni-NTA resin. Glutathione agarose resin was incubated in batch with the cleared lysate in the presence of 0.1 % (v/v) Tween 20 for 1 h at 4 ºC, washed with wash buffer C (25 mM HEPES, 250 mM NaCl, 0.1 % Tween 20, 1 mM DTT, 2 mM benzamidine-HCl, pH 8.0), and proteins were eluted on a gravity column with elution buffer C (50 mM HEPES, 150 mM NaCl, 10 mM reduced L-glutathione, 1 mM DTT, 2 mM benzamidine-HCl, pH 8.0). Fractions containing the proteins were pooled and further purified by size exclusion chromatography using a Superose 6 10/300 column (GE Healthcare) equilibrated with storage buffer. Glycerol and DTT were added to final concentrations of 10 % (v/v) and 1 mM, respectively, and aliquots were flash-frozen in liquid nitrogen and stored at -80 ºC.

### GST pull-down

50 pmol of purified GST::SMK-1(146-282)::6xHis were incubated with 50 pmol of purified 6xHis::MBP::HCP-4::StrepTagII for 1 h at 4 ºC in 150 μL pull-down buffer (50 mM HEPES, 100 mM NaCl, 5 mM DTT, pH 7.5) containing 0.1 % Tween 20 and supplemented with 15 μL of glutathione agarose resin. After washing the resin with 3 × 500 μL pull-down buffer, proteins were eluted with 50 µL pull-down buffer containing 15 mM reduced L-glutathione. Eluted proteins were separated by SDS-PAGE and visualized by Coomassie Blue staining.

### Immunoblotting

For immunoblots of *C. elegans* lysate, ~60 adult hermaphrodites were collected into 500 μL M9 buffer and washed 2 × with M9 buffer and 2 × with M9 containing 0.05 % Triton X-100. To 100 μL of worm suspension, 33 μL 4x SDS-PAGE sample buffer [250 mM Tris-HCl, pH 6.8, 30 % (v/v) glycerol, 8 % (w/v) SDS, 200 mM DTT and 0.04 % (w/v) bromophenol blue] and ~20 μL of glass beads were added. Samples were incubated for 3 min at 95 ºC and vortexed for 2 × 5 min with intermittent heating. After centrifugation at 20000 × g for 1 min at room temperature, proteins in the supernatant were resolved by 10 % SDS-PAGE and transferred to a 0.2-μm nitrocellulose membrane (Hybond ECL, Amersham Pharmacia Biotech). Membranes were rinsed 3 × with TBS (50 mM Tris-HCl, 145 mM NaCl, pH 7.6), blocked with 5 % non-fat dry milk in TBST (TBS containing 0.1 % Tween 20), and probed at 4 ºC overnight with mouse monoclonal anti-FLAG M2 (1:1000, Sigma-Aldrich) and mouse monoclonal anti-α-tubulin B512 (1:5000, Sigma-Aldrich). The membrane was washed 3 × with TBST, incubated with goat polyclonal anti-mouse IgG antibody coupled to HRP (1:10000; Jackson ImmunoResearch 115-035-044) for 1 hour at room temperature, and washed again 3 × with TBST. Proteins were detected by chemiluminescence using Pierce ECL Western Blotting Substrate (Thermo Scientific) and X-ray film (Amersham, GE Healthcare).

### Statistical analysis

Statistical analysis was performed with Prism 9.0 software (GraphPad). Statistical significance was determined by ANOVA on ranks (Kruskal-Wallis nonparametric test) followed by Dunn’s multiple comparison test, or by a two-sided Mann-Whitney test, where \*\*\*\**P* < 0.0001, \*\*\**P* < 0.001, \*\**P* < 0.01, \**P* < 0.05, and *ns* = not significant, *P* > 0.05. The analytical method used is specified in the figure legends.

## ACKNOLEDGEMENTS

The authors acknowledge the Biochemical and Biophysical Technologies Scientific Platform at i3S for support. Some strains were provided by the *Caenorhabditis* Genetics Center (CGC), which is funded by NIH Office of Research Infrastructure Programs (P40 OD010440). Funding for this project was provided by the European Research Council under the European Union’s Seventh Framework Programme (ERC grant agreement n ° ERC-2013-StG-338410-DYNEINOME). R.G. is supported by FCT Principal Investigator position CEECIND/00333/2017 and H.R. was supported by FCT PhD fellowship SFRH/BD/103495/2014. The authors declare no competing financial interests.

## FIGURE LEGENDS

**Figure S1.**
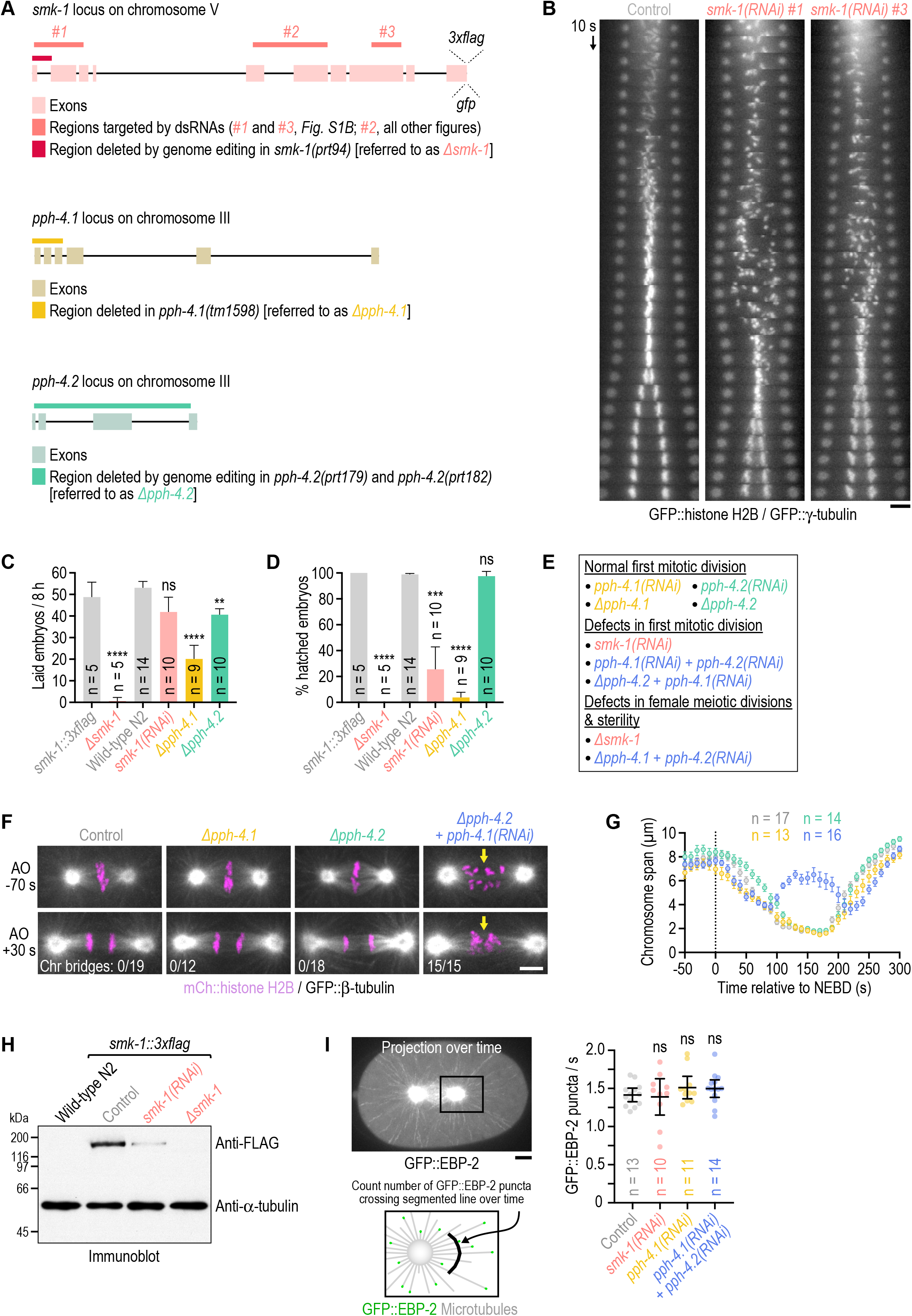
Additional characterization of PP4 inhibition. **(A)** Genomic loci of PP4 subunits with annotation of existing mutations (*∆pph-4*.*1*) and modifications introduced by genome-editing for this study (*smk-1::3xflag, smk-1::3xgfp, ∆smk-1, ∆pph-4*.*2*). Note that the *∆smk-1* mutation was introduced into the *smk-1::3xflag* background. Regions in *smk-1* targeted by dsRNAs are also indicated. **(B)** Time-aligned kymographs generated from time-lapse movies of one-cell embryos co-expressing GFP::histone H2B (HIS-58) and GFP::γ-tubulin (TBG-1). Of the three dsRNAs described in *(A)*, #1 and #3 were used here, and #2 was used for all other experiments. Scale bar, 5 µm. **(C), (D)** Number of embryo progeny laid by a single mother in an 8-h interval *(C)*, and embryonic viability *(D)*, plotted as the percentage of hatched embryo progeny from a single mother (mean of *n* mothers ± 95 % CI). Mothers were homozygous for the mutations analyzed. The *smk-1::3xflag* strain serves as the control for *∆smk-1*, and the wild-type N2 strain serves as the control for the other conditions. Statistical significance (control versus perturbations) was determined by ANOVA on ranks (Kruskal-Wallis nonparametric test) followed by Dunn’s multiple comparison test. \*\*\*\**P* < 0.0001; \*\*\**P* < 0.001; \*\**P* < 0.01; *ns* = not significant, *P* > 0.05. **(E)** Summary of the effect of different PP4 inhibition conditions on chromosome segregation in embryonic mitosis and female meiosis. Homozygous *∆smk-1* animals and homozygous *∆pph-4*.*1* animals injected with dsRNA against *pph-4*.*2* only produce a few embryos before becoming sterile and could therefore not be used for characterization of mitotic PP4 function. **(F)** Selected images from time-lapse movies of one-cell embryos co-expressing mCherry::histone H2B (HIS-11) and GFP::β-tubulin (TBB-2). Time is relative to anaphase onset (AO). Arrows highlight defective chromosome congression and segregation in PP4-inhibited embryos. The number of anaphases with chromatin (chr) bridges relative to the total number of anaphases examined is indicated. Scale bar, 5 µm. **(G)** Chromosome span (mean of *n* embryos ± SEM) versus time relative to NEBD, measured in one-cell embryos such as those shown in *(F)*. Conditions are color-coded as in *(F)*. **(H)** Anti-FLAG immunoblot of adult animals, showing expression levels of endogenous 3xFLAG-tagged SMK-1, the absence of SMK-1::3xFLAG signal in the null mutant *∆smk-1*, and residual SMK-1::3xFLAG signal after RNAi-mediated depletion. Anti-α-tubulin antibody serves as the loading control. Molecular weight is indicated in kilodaltons (kDa). **(I)** *(left top)* Maximum intensity projection over successive frames from a time-lapse movie of a one-cell embryo expressing the microtubule plus end marker GFP::EBP-2. Scale bar, 5 µm. *(left bottom)* Schematic illustrating the assay for quantification of microtubule nucleation rate. *(right)* Rate at which GFP::EBP-2 particles cross a segmented line near centrosomes (mean of *n* embryos ± 95 % CI) as a measure of microtubule nucleation rate. Statistical significance (control versus perturbations) was determined by ANOVA on ranks (Kruskal-Wallis nonparametric test) followed by Dunn’s multiple comparison test. *ns* = not significant, *P* > 0.05.

**Figure S2.**
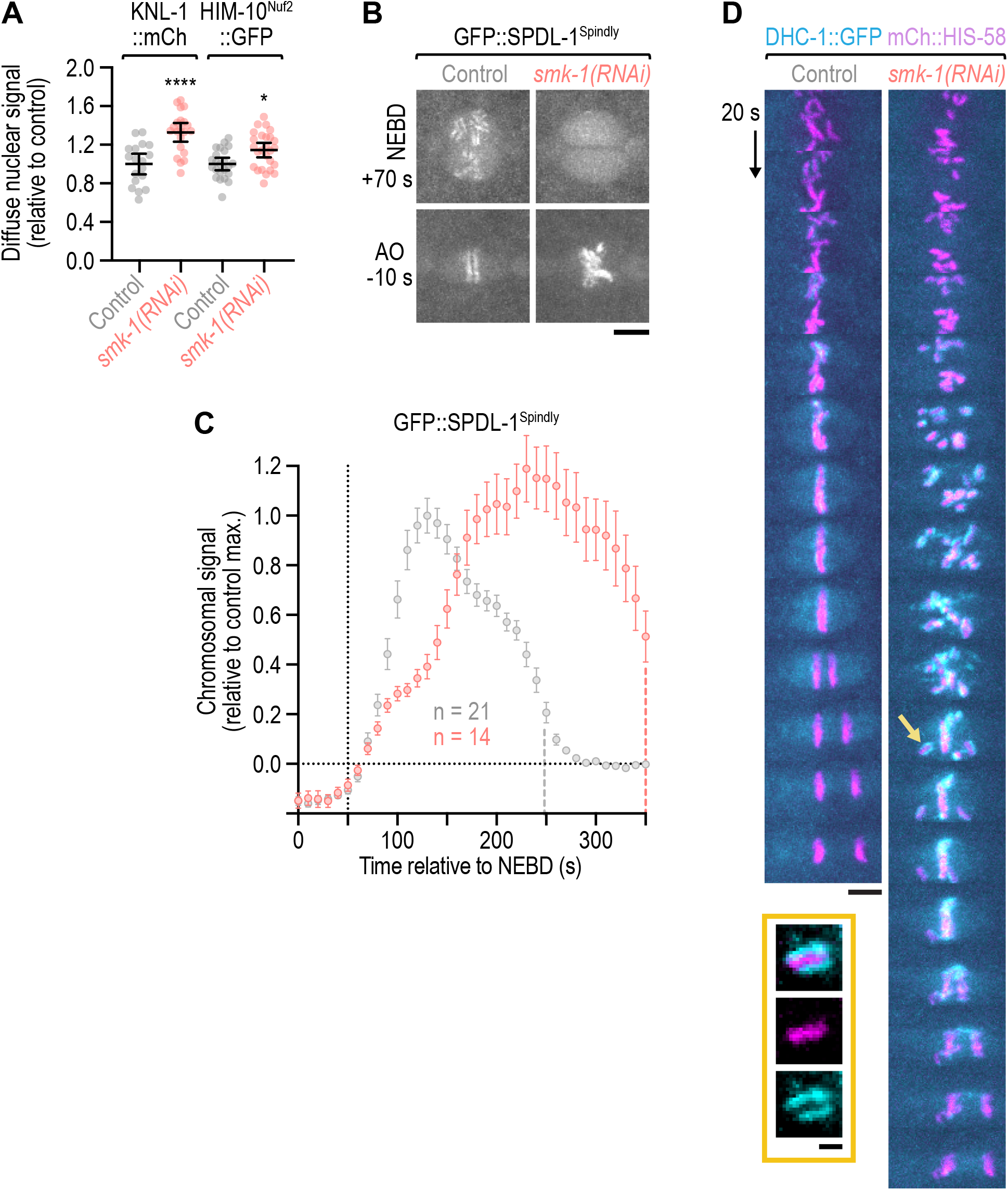
Nuclear import of outer kinetochore components is unaffected by PP4 inhibition; the outermost kinetochore component dynein hyperaccumulates in late prometaphase following PP4 inhibition. **(A)** Intensity of the non-chromosomal (diffuse) nuclear signal of fluorescently tagged outer kinetochore components at NEBD in the one-cell embryo (mean ± 95 % CI), normalized to the mean of the respective control. Statistical significance [control versus *smk-1(RNAi)*] was determined by the Mann-Whitney test. \*\*\*\**P* < 0.0001; \**P* < 0.05. **(B)** Selected images from time-lapse movies of one-cell embryos co-expressing transgenic GFP::SPDL-1 in an *spdl-1* knock-out background and mCherry-tagged histone H2B (HIS-58). Only the GFP::SPDL-1 signal is shown. AO, anaphase onset. Scale bar, 5 µm. **(C)** Intensity of the chromosomal GFP::SPDL-1 signal (mean of *n* embryos ± SEM) versus time relative to NEBD, determined by averaging the signal of the top 15 local maxima detected on chromosomes in time-lapse movies of one-cell embryos. Traces are normalized to the peak signal in the control. Vertical dashed lines mark the average time of anaphase onset. **(D)** Selected images from time-lapse movies of one-cell embryos co-expressing endogenously GFP-tagged dynein heavy chain (DHC-1) and transgenic mCherry::histone H2B (HIS-58). Image sequences start at the same time point after NEBD. Arrow points to the unaligned chromosome shown at greater magnification. Scale bar, 5 µm; magnified chromosome, 2 µm.

**Figure S3.**
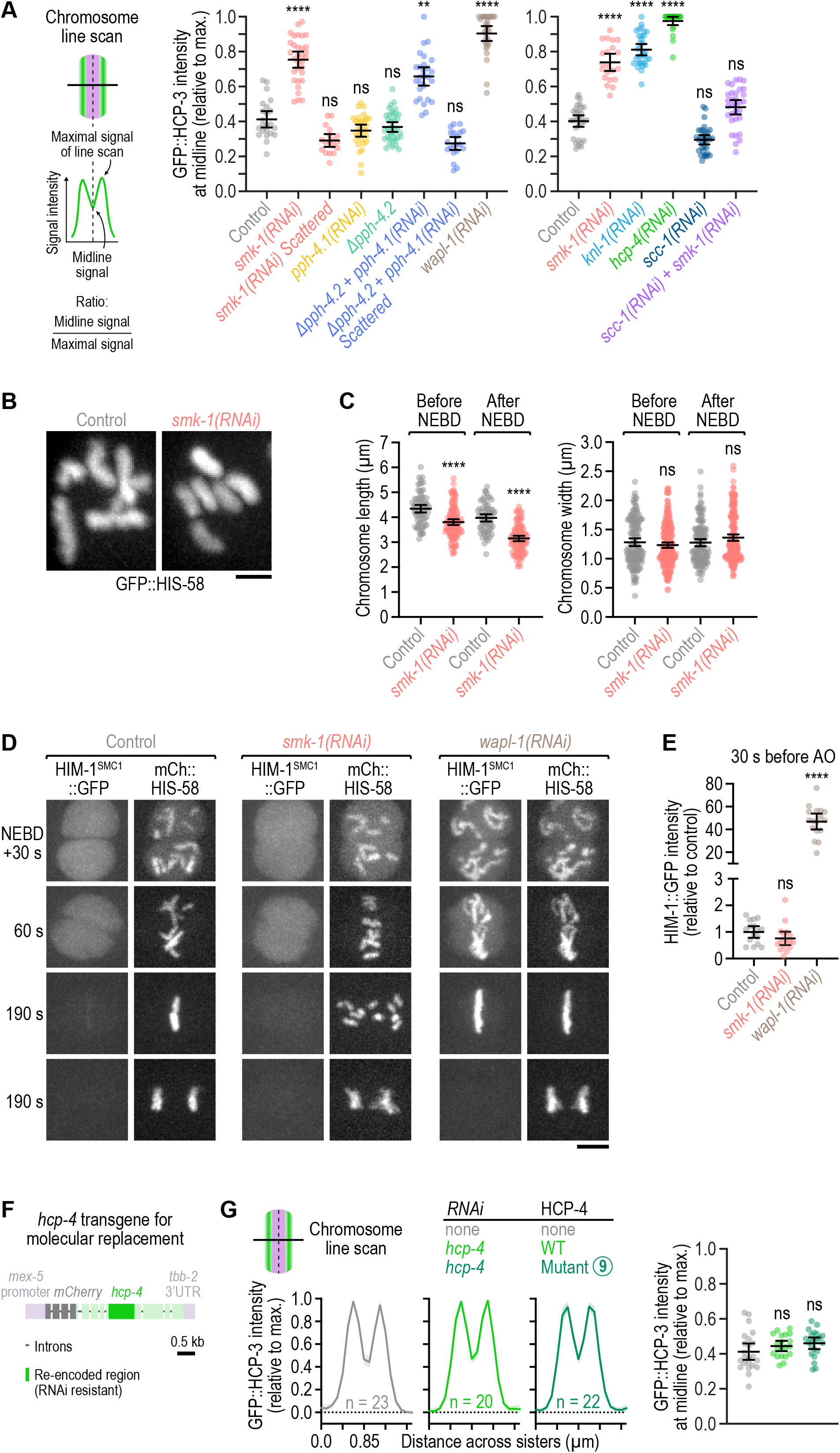
Quantification of the sister centromere resolution defects shown in *Fig. 3*; mitotic chromosomes have a compact morphology and do not exhibit defects in cohesin removal following PP4 inhibition; the SMK-1–HCP-4^CENP-C^ interaction is dispensable for sister centromere resolution. **(A)** Quantification of the line scans shown in *Fig. 3D* and *F* (left and right graph, respectively). GFP::HCP-3 signal intensity at the chromosome midline relative to the maximal signal intensity in the line scan (mean of *n* chromosomes from at least 10 embryos ± 95 % CI) was determined as illustrated in the schematic. Statistical significance (control versus perturbations) was determined by ANOVA on ranks (Kruskal-Wallis nonparametric test) followed by Dunn’s multiple comparison test. \*\*\*\**P* < 0.0001; \*\*\**P* < 0.001; \*\**P* < 0.01; *ns* = not significant, *P* > 0.05. **(B)** Morphology of early prometaphase chromosomes in one-cell embryos, marked by GFP::histone H2B (HIS-58). Scale bar, 2 µm. **(C)** Length and width of mitotic chromosomes in late prophase and early prometaphase (mean ± 95 % CI), derived from 3D measurements in z-stacks acquired of GFP-marked chromosomes as shown in *(B)*. Data points correspond to values of individual chromosomes from at least 5 embryos per condition, normalized to the mean of the control. Statistical significance [control versus *smk-1(RNAi)*] was determined by the Mann-Whitney test. \*\*\*\**P* < 0.0001; *ns* = not significant, *P* > 0.05. **(D)** Selected images from time-lapse movies of one-cell embryos co-expressing endogenously tagged HIM-1::GFP (the SMC1 subunit of the cohesin complex) and transgenic mCherry::histone H2B (HIS-58). Time is relative to NEBD. Scale bar, 5 µm. **(E)** Integrated intensity of chromosomal HIM-1::GFP signal (mean ± 95 % CI) three frames before anaphase onset (AO) in the one-cell embryo, normalized to the mean of the control. Signal was measured in embryos such as those shown in *(D)*. Statistical significance (control versus perturbations) was determined by ANOVA on ranks (Kruskal-Wallis nonparametric test) followed by Dunn’s multiple comparison test. \*\*\*\**P* < 0.0001; *ns* = not significant, *P* > 0.05. **(F)** Schematic of the mCherry-tagged RNAi-resistant HCP-4^CENP-C^ transgene used for rescue experiments in embryos. **(G)** *(left)* Line scan profiles (mean of *n* chromosomes from at least 10 embryos ± 95 % CI), generated as described in *Fig. 3C. (right)* Ratio of GFP::HCP-3 signal intensity at the midline of condensed chromosomes to the maximal signal intensity in the line scan (mean of *n* chromosomes from at least 10 embryos ± 95 % CI), determined as described in *(A)*. Statistical significance (control versus perturbations) was determined by ANOVA on ranks (Kruskal-Wallis nonparametric test) followed by Dunn’s multiple comparison test. *ns* = not significant, *P* > 0.05.

## SUPPLEMENTAL MOVIE LEGENDS

**Movie S1**.

One-cell embryos co-expressing GFP::histone H2B and GFP::γ-tubulin. Control *(left)* and *smk-1(RNAi) (right)*, aligned relative to NEBD. Time-lapse is 2 s and playback speed is 12 frames per second.

**Movie S2**.

One-cell embryos co-expressing GFP::β-tubulin (white) and mCherry::histone H2B (magenta). Control *(left)* and *smk-1(RNAi) (right)*, aligned relative to NEBD. Time-lapse is 10 s and playback speed is 6 frames per second.

**Movie S3**.

One-cell embryos co-expressing GFP::β-tubulin (white) and mCherry::histone H2B (magenta). *∆pph-4*.*1 (top left), ∆pph-4*.*2 (top right), pph-4*.*1(RNAi) (bottom left)*, and *∆pph-4*.*2* + *pph-4*.*1(RNAi) (bottom right)*, aligned relative to NEBD. Time-lapse is 10 s and playback speed is 6 frames per second.

**Movie S4**.

One-cell embryos co-expressing GFP::histone H2B and GFP::γ-tubulin. *klp-19(RNAi) (left)* and *klp-19(RNAi);smk-1(RNAi) (right)*, aligned relative to NEBD. Time-lapse is 10 s and playback speed is 6 frames per second.

**Movie S5**.

One-cell embryos expressing GFP::HCP-4^CENP-C^. Control *(left)* and *smk-1(RNAi) (right)*, aligned relative to NEBD. Time-lapse is 10 s and playback speed is 6 frames per second.

**Movie S6**.

One-cell embryos co-expressing HIM-10^Nuf2^::GFP and mCherry::histone H2B. Control *(left)* and *smk-1(RNAi) (right)*, aligned relative to NEBD. Time-lapse is 10 s and playback speed is 6 frames per second.

**Movie S7**.

One-cell embryos co-expressing dynein heavy chain DHC-1::GFP (cyan) and mCherry::histone H2B (red). Control *(left)* and *smk-1(RNAi) (right)*, aligned relative to NEBD. Time-lapse is 10 s and playback speed is 6 frames per second.

**Movie S8**.

One-cell embryo co-expressing SMK-1::GFP and mCherry::histone H2B. Time-lapse is 10 s and playback speed is 6 frames per second.

**Movie S9**.

One-cell embryos co-expressing GFP::HCP-3^CENP-A^ and mCherry::histone H2B. Control *(left)* and *smk-1(RNAi) (right)*, aligned relative to NEBD. Time-lapse is 10 s and playback speed is 6 frames per second.

**Movies S10**.

One-cell embryos expressing GFP::HCP-3^CENP-A^. Control *(left)* and *knl-1(RNAi) (right)*, aligned relative to NEBD. Time-lapse is 10 s and playback speed is 6 frames per second.

**Movie S11**.

One-cell embryos co-expressing GFP::histone H2B and γ-tubulin::GFP. *smk-1(RNAi) (left)* and *∆ska-1;smk-1(RNAi) (right)*, aligned relative to NEBD. Time-lapse is 10 s and playback speed is 6 frames per second.

**Movie S12**.

One-cell embryos co-expressing GFP::histone H2B and γ-tubulin::GFP. *∆ska-1;smk-1(RNAi);C09B9*.*4(RNAi) (left)* and *∆ska-1;smk-1(RNAi);rod-1(RNAi) (right)*, aligned relative to NEBD. *C09B9*.*4(RNAi)* serves as the control for double RNAi. Time-lapse is 10 s and playback speed is 6 frames per second.

**Table S1:**
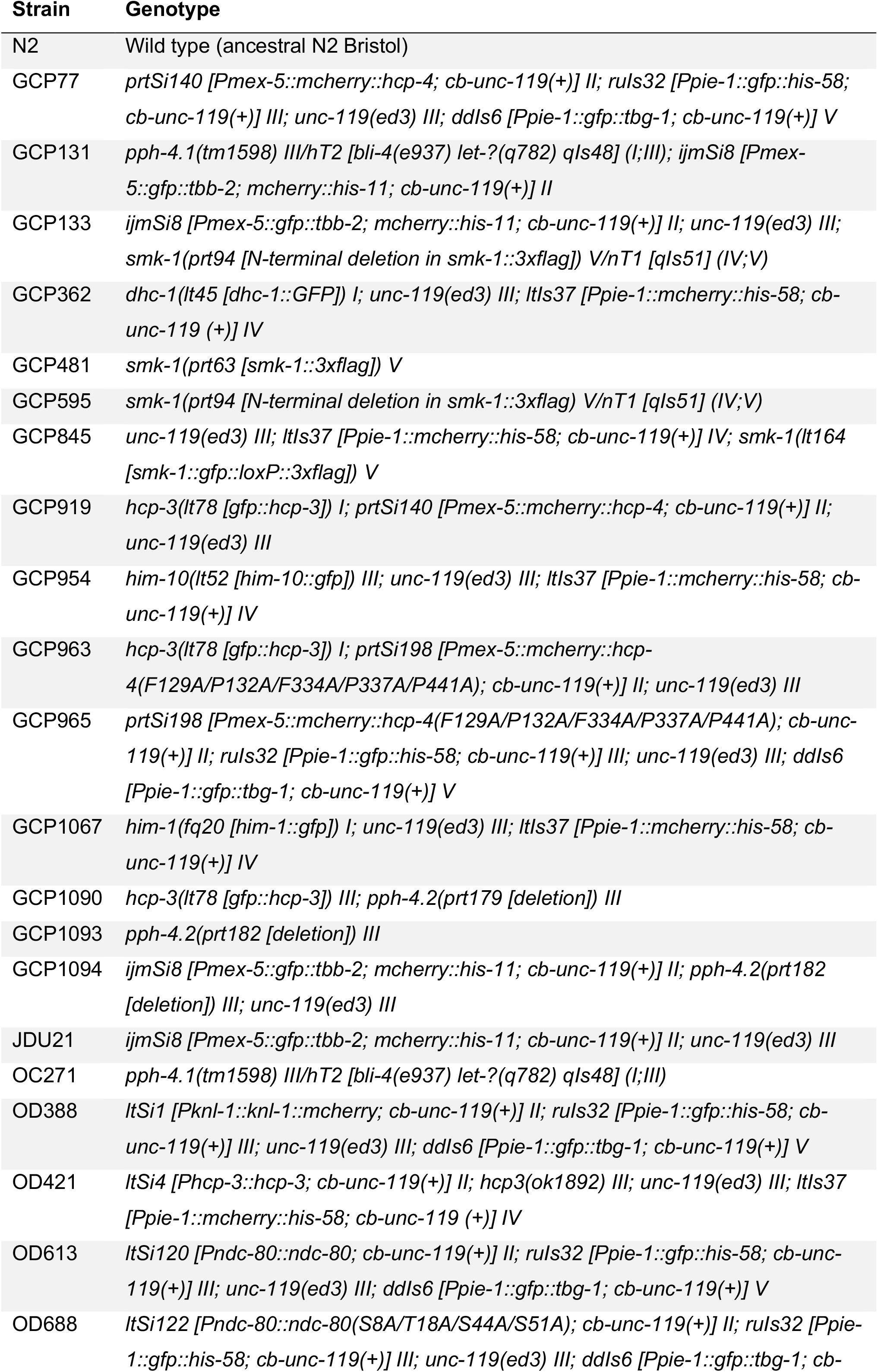
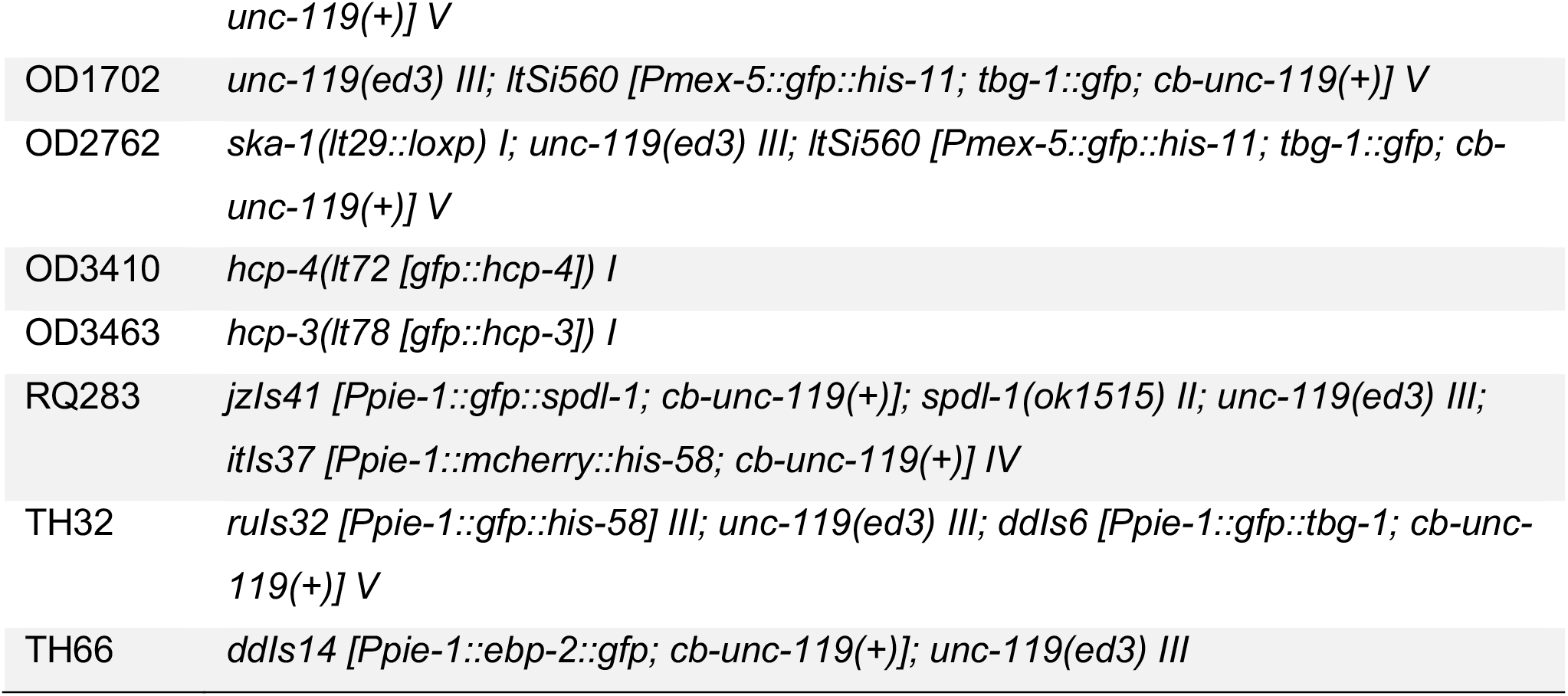
*C. elegans* strains.

**Table S2:**
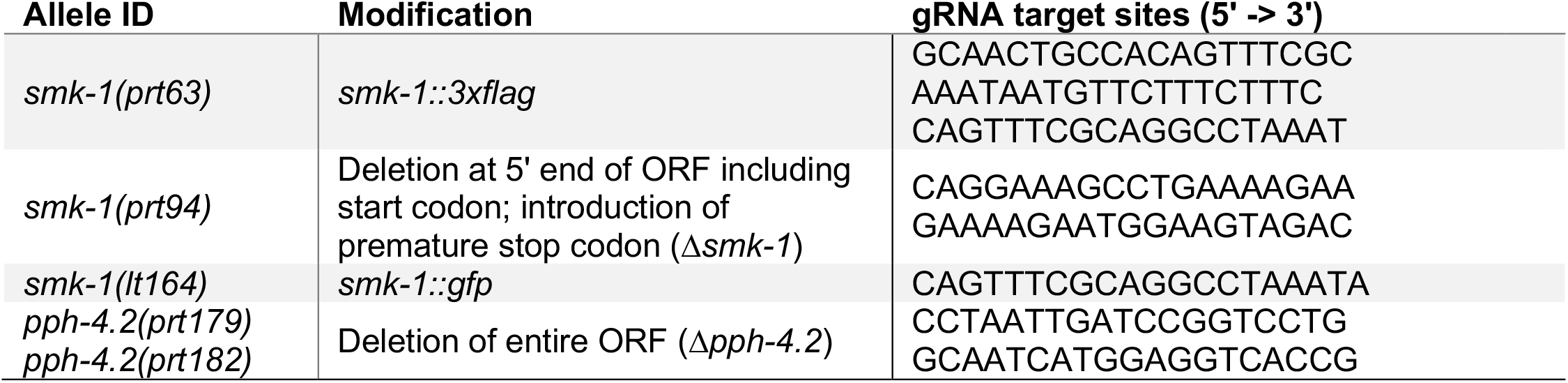
Genomic sequences targeted by gRNAs for CRISPR/Cas9-mediated genome editing.

**Table S3:**
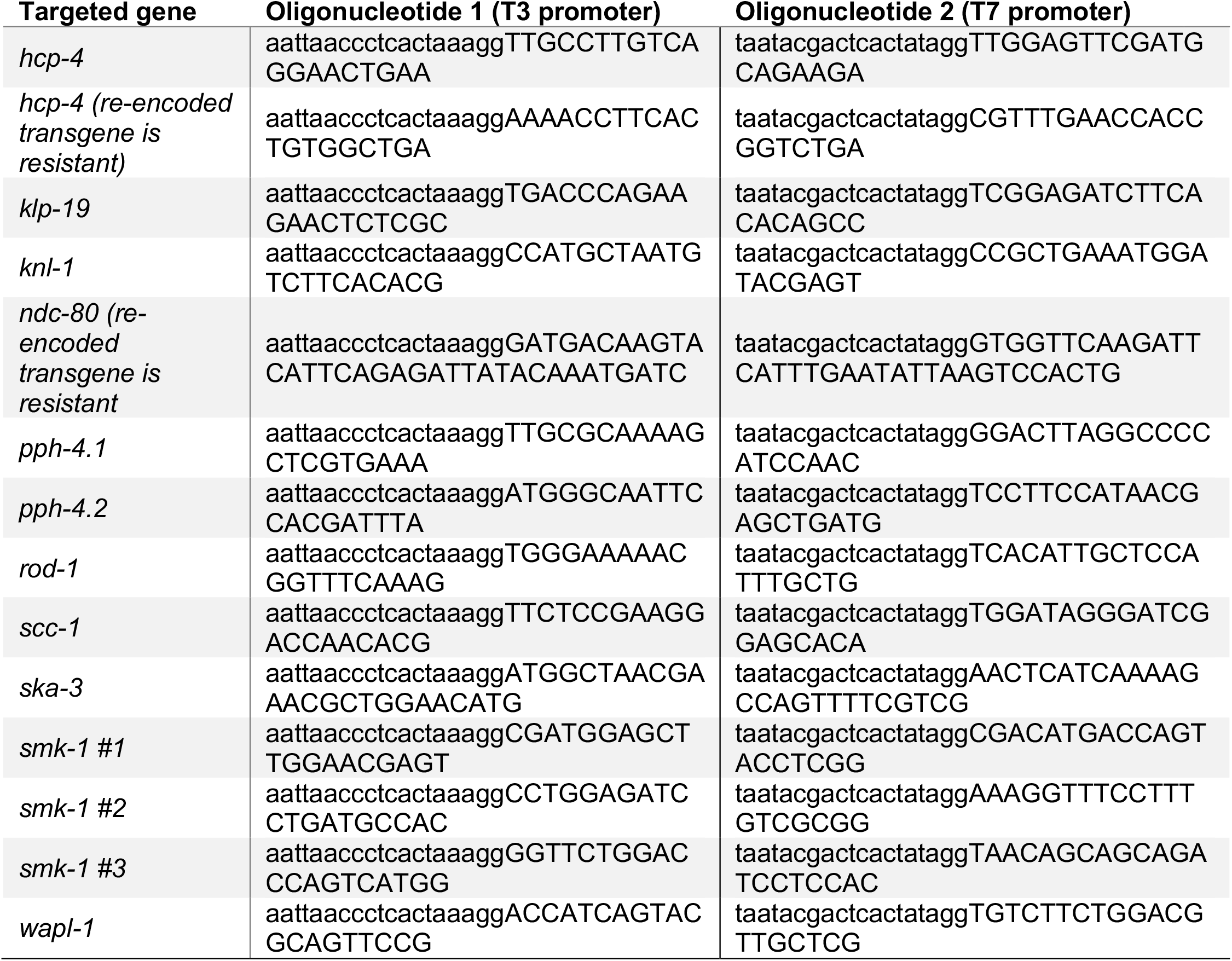
Oligonucleotides for dsRNA production.

